# Transcriptomic and morphophysiological evidence for a specialized human cortical GABAergic cell type

**DOI:** 10.1101/216085

**Authors:** Eszter Boldog, Trygve Bakken, Rebecca D. Hodge, Mark Novotny, Brian D. Aevermann, Judith Baka, Sándor Bordé, Jennie L. Close, Francisco Diez-Fuertes, Song-Lin Ding, Nóra Faragó, Ágnes K. Kocsis, Balázs Kovács, Jamison M. McCorrison, Jeremy A. Miller, Gábor Molnár, Gáspár Oláh, Attila Ozsvár, Márton Rózsa, Soraya Shehata, Kimberly A. Smith, Susan M. Sunkin, Danny N. Tran, Pratap Venepally, Abby Wall, László G. Puskás, Pál Barzó, Frank J. Steemers, Nicholas J. Schork, Richard H. Scheuermann, Roger S. Lasken, Ed S. Lein, Gábor Tamás

**Affiliations:** MTA-SZTE Research Group for Cortical Microcircuits, Department of Anatomy, Physiology and Neuroscience, University of Szeged, Közép fasor 52., Szeged, H-6726, Hungary; Allen Institute for Brain Science, 615 Westlake Avenue North, Seattle, WA 98109, USA; J. Craig Venter Institute, 4120 Capricorn Lane, La Jolla, CA 92037, USA; Illumina, Inc., 5200 Illumina Way, San Diego, CA 92122 USA; Laboratory of Functional Genomics, Department of Genetics, Biological Research Center, Hungarian Academy of Sciences, Temesvári krt. 62, H-6726, Szeged, Hungary; Department of Neurosurgery, University of Szeged, Hungary, Semmelweis u. 6., Szeged, H-6725 Hungary; Department of Pathology, 9500 Gilman Drive, University of California, San Diego, CA 92093 USA

**Author notes:** These authors have equal contributions (Eszter Boldog, Trygve Bakken, Rebecca D. Hodge). Correspondence should be addressed to Ed S. Lein and Gábor Tamás.

**Keywords:** human, neocortex, interneuron, layer 1, cell type, transcriptomics, microcircuit

## Abstract

We describe convergent evidence from transcriptomics, morphology and physiology for a specialized GABAergic neuron subtype in human cortex. Using unbiased single nucleus RNA sequencing, we identify ten GABAergic interneuron subtypes with combinatorial gene signatures in human cortical layer 1 and characterize a novel group of human interneurons with anatomical features never described in rodents having large, “rosehip”-like axonal boutons and compact arborization. These rosehip cells show an immunohistochemical profile (GAD1/CCK-positive, CNR1/SST/CALB2/PVALB-negative) matching a single transcriptomically-defined cell type whose molecular signature is not seen in mouse cortex. Rosehip cells make homotypic gap junctions, predominantly target apical dendritic shafts of layer 3 pyramidal neurons and inhibit backpropagating pyramidal action potentials in microdomains of the dendritic tuft. These cells are therefore positioned for potent local control of distal dendritic computation in cortical pyramidal neurons.

---

Understanding the cellular and circuit organization of the neocortex, the substrate for much of higher cognitive function, has been a topic of intense study since the seminal work of Ramón y Cajal ^1^. Morpho-physiological characterization using in vitro slice physiology has been the standard for decades ^2^, but this approach suffers from undersampling, difficulties in quantitative classification of cell types ^3^, and applicability largely to model organisms at sufficient scale to seriously handle neuronal diversity. Recent advances in single cell transcriptomics has offered a new approach for unbiased, high-throughput quantitative surveys of molecularly defined cell types ^4–6^ that is in principle applicable to tissues from any species including human. Initial application to mouse cortex has revealed approximately 50 transcriptomic types, demonstrating both the feasibility of the approach and the complexity of the system. There is now great promise in combining these traditional (morpho-electric) and nouveau (transcriptomic) approaches for an unbiased molecular classification and then characterization of these types.

Recent systematic efforts in rodent have provided insight into the cellular composition and organization of rodent neocortical circuits, suggesting the presence of a several dozen inhibitory and excitatory cell types ^3–5,7^. However, conservation of cellular and circuit principles in human cortex is largely assumed but largely untested to date. Indeed there is evidence for significant neuronal differences between rodents and human; for example, emerging results suggest that distinct membrane ^8,9^ and synaptic ^10–13^ properties and dendritic complexity ^14–16^ of human neurons might contribute to human specific signal processing. With the increasing study of mouse cortex as a model for understanding human cognition it is essential to establish whether the cellular architecture of human is conserved or whether there are specialized cell types and system properties that cannot be modeled in rodents. Here we combine single nucleus transcriptomics and in vitro slice physiology to study GABAergic neurons in layer 1 of human cortex and provide multiple convergent lines of evidence for the identification of a cell type with human specialized features.

## Results

To allow an unbiased survey of transcriptionally-defined cell types in human cortical tissue we adapted methods for single nucleus RNA sequencing ^17,18^ to profile large numbers of nuclei from frozen postmortem brain specimens (Fig. 1A). Briefly this method involved microdissection of regions of interest from fluorescent Nissl-stained vibratome sections of cortex, tissue homogenization to liberate nuclei, NeuN staining and FACS isolation, and Smart-seq2 based library preparation ^19^. We applied this strategy to profile n=769 quality control passed NeuN-positive neurons and n=102 NeuN-negative non-neuronal cells across 2 individuals from microdissected layer 1 of the middle temporal gyrus, expected to predominantly contain inhibitory neurons. Iterative clustering was used to group nuclei with similar transcriptional profiles, thereby identifying a robust set of transcriptomic-defined cell types (Fig. 1B). Based on expression of known marker genes (Suppl. Fig. 1A), clusters corresponded to all major classes of neural cell types that were expected to be captured. These included major non-neuronal cell types (microglia, astrocytes, oligodendrocyte precursor cells (OPCs) and oligodendrocytes) and one excitatory neuron type sampled from upper cortical layer 2 incidentally included in the layer 1 dissection (Fig. 1C). In addition, eleven clusters corresponding to GABAergic neuron subtypes were identified (numbered by relative abundance), including three highly distinctive clusters (i1, i2, i5) and a larger set of more closely-related cell types.

**Figure 1.**
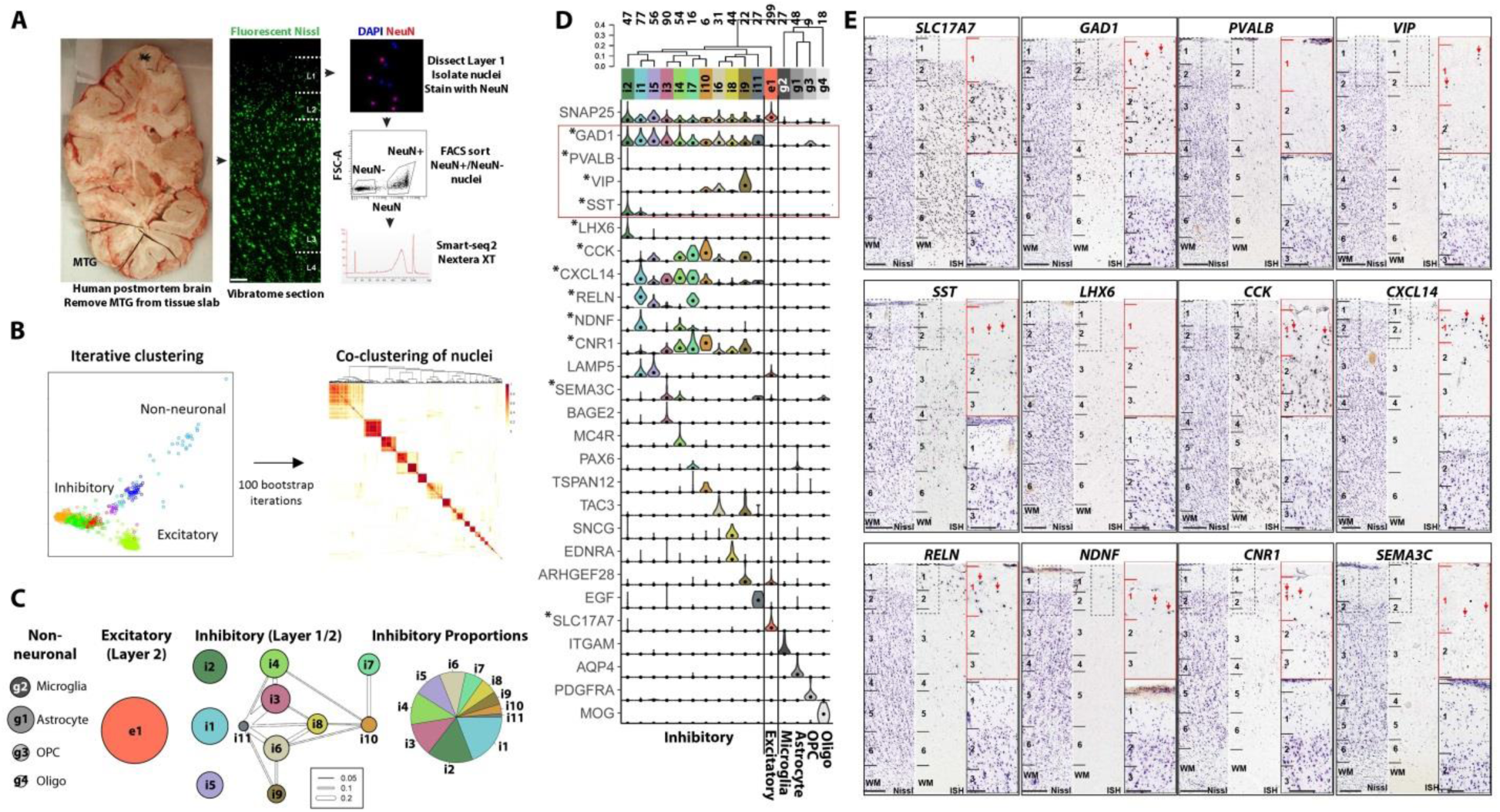
Identification of transcriptomic cell types in Layer 1 of human temporal cortex. **A**, Isolation of single nuclei from post-mortem adult human cortex for RNA-sequencing. Scale bar=50 µm. **B**, Left: Nuclei were grouped based on similar gene expression profiles using a fully automated iterative clustering procedure. Right: Cluster robustness was assessed by randomly subsampling 80% of nuclei and reclustering. Red boxes along diagonal and white off-diagonal demonstrate consistent, well separated clusters. **C**, 4 non-neuronal, 1 excitatory and 11 inhibitory neuron clusters were identified, although the excitatory cluster and one inhibitory cluster were likely in layer 2 incidentally captured due to incidental capture with layer 1 dissection. For each cluster, the constellation diagram shows the broad cell class membership (based on canonical marker gene expression), relative size (disc area), and discreteness (line thickness proportional to average co-clustering) of clusters. **D**, Clusters arranged by transcriptomic similarity based on hierarchical clustering, with the expression distributions of selective marker genes shown across clusters as violin plots. Numbers denote the number of cells in each cluster. **E**, ISH of select marker genes in human temporal cortex at low magnification (left columns with near adjacent Nissl stain for cytoarchitectonic laminar identification) and high magnification in layers 1-3 (right column). Red arrows highlight cells expressing genes in layer 1. Note that *LHX6* marks a single cluster (i2) that is not expressed in Layer 1 and therefore nuclei in this cluster were likely sampled from upper Layer 2. Other clusters are restricted to layer 1 (e.g. *NDNF*+) or may be distributed across Layers 1 and 2. Scale bars=250 µm (low mag), 100 µm (high mag).

Transcriptomic cell types displayed highly selective gene expression (Fig. 1D, Suppl. Fig. 1A). For example, the pan-neuronal gene synaptosomal-associated protein 25 (*SNAP25*) clearly differentiated neuronal from non-neuronal types, which were in turn differentiated by highly specific marker genes. Glutamic acid decarboxylase 1 (*GAD1*) clearly delineated the GABAergic neurons. In cortical layers 2-6, most GABAergic neurons have mutually exclusive expression of parvalbumin (*PVALB*), somatostatin (*SST*) or vasoactive intestinal peptide (*VIP*) ^20^. In contrast, *Pvalb* and *Sst* are not expressed in mouse layer 1 by *in situ* hybridization (ISH), while *Vip* labels only sparse cell populations (Suppl. Fig. 1B). Interestingly, both *SST* and *VIP* (but not *PVALB*) are seen in human MTG layer 1 by ISH (Fig. 1E). The layer 1 MTG transcriptomic clusters expressed either *SST* (i1,i2), *VIP* (i6,i9,i10), or neither marker, although cluster i2 represents a cell type restricted to layer 2 since it also expresses *LHX6* which is not found in layer 1 (Fig. 1D,E). Therefore, there appear to be ten inhibitory cell types within layer 1, although it is not clear whether any of these types are completely restricted to layer 1. These clusters in layer 1 express different combinations of known markers of layer 1 interneurons, including cholecystokinin (*CCK*), reelin (*RELN*), neuron derived neurotrophic factor (*NDNF*), and lysosomal associated membrane protein family member 5 (*LAMP*), many of which were confirmed to have expression in layer 1 by ISH (Fig. 1E). Furthermore, each cluster showed highly selective expression of known and previously uncharacterized individual marker genes. Interestingly, given the proximity of layer 1 to the overlaying pia, several of these markers appear to be related to interaction with endothelia, including endothelin receptor type A (*EDNRA*) and epidermal growth factor (*EGF*). To summarize, this unbiased transcriptomic approach identified ten GABAergic interneuron subtypes in layer 1 that have distinctive combinatorial and specific gene expression signatures suggestive of distinct morphological and functional properties.

### Rosehip cells: novel morphological features in layer 1 of the human cerebral cortex

In parallel to the transcriptomic approach we developed an initial database of whole cell recorded, biocytin filled interneurons in layer 1 of 350 µm thick slices of nonpathological human samples prepared from surgically removed pieces of parietal, frontal and temporal cortices ^10,11,21^. Initially, there was no preference when approaching particular cell types during recordings, apart from having the soma of the patch clamped cells in layer 1. This unbiased approach yielded interneurons with complete somatodendritic and axonal morphological recovery (n=76). Light microscopic examination of each cell identified neurons with previously described morphological features, e.g. neurogliaform cells (n=16, 21%; Fig. 2D) ^20,22,23^.

**Figure 2.**
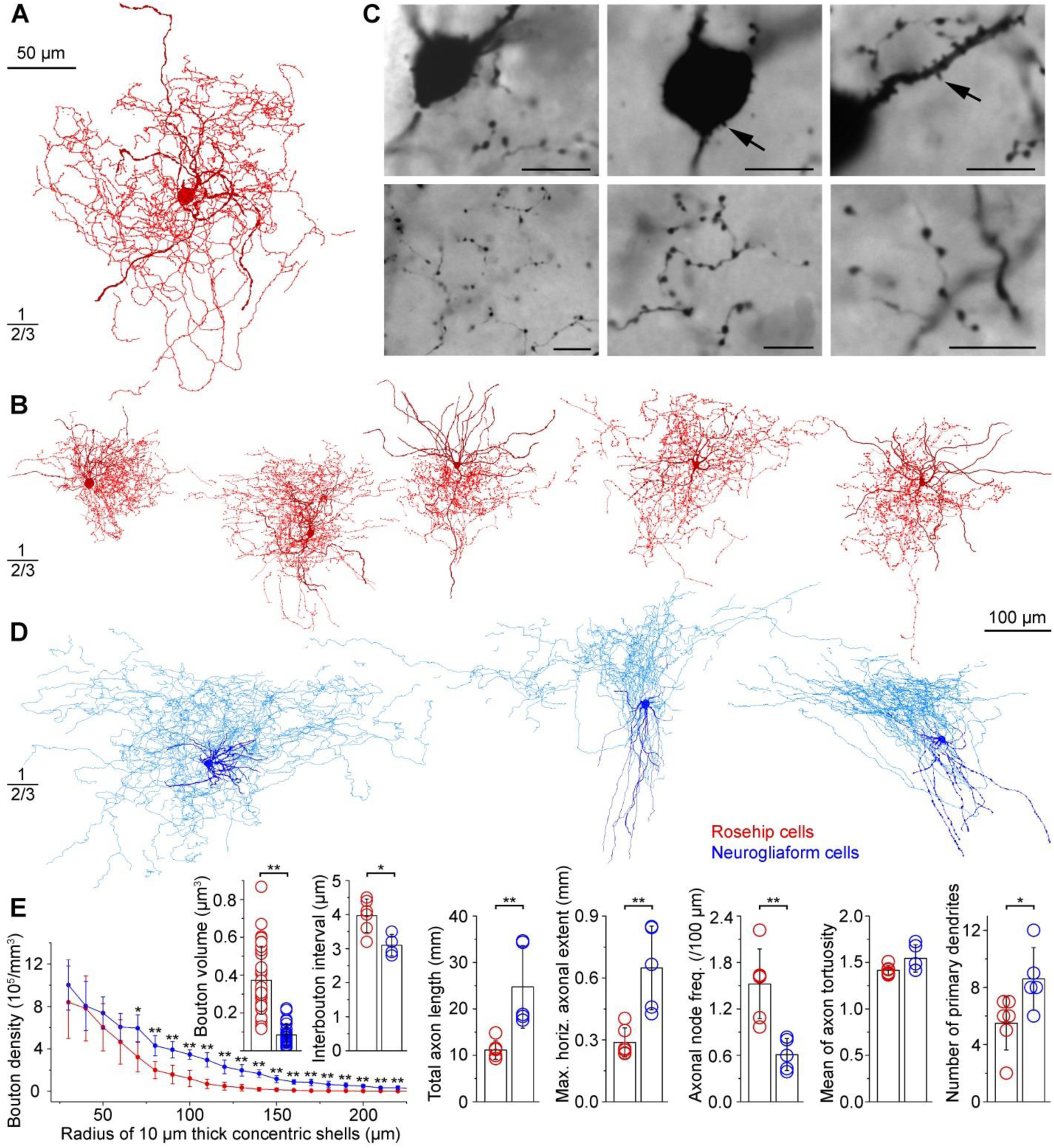
Morphological phenotype of rosehip cells in layer 1 of the human cerebral cortex. **A, B,** Anatomical reconstructions of rosehip cells biocytin filled during whole cell recordings (somata and dendrites, burgundy; axons, red). **C,** Top, Light micrographs of rosehip cells showing somata and proximal dendrites with stub-like spines (arrows). Bottom, Axons of rosehip cells arborized densely with large, round boutons. Scale bars: 10 µm. **D,** Anatomical reconstructions of neurogliaform cells in layer 1 of the human cerebral cortex (somata and dendrites, dark blue; axons, light blue). **E**, Quantitative comparison of axonal and dendritic parameters of rosehip (red) and neurogliaform (blue) cells. Left, bouton densities determined by Sholl analysis in 10 µm thick spherical shells were higher in neurogliaform cells 70-220 µm from the soma. Right, Numbers of primary dendrites, total axonal length and maximal horizontal extent of the axon of neurogliaform cells exceeded that of rosehip cells. Axonal tortuosity was similar in the two cell types, however, the frequency of axonal branch points in rosehip cells was 2.5 times that of neurogliaform cells. However, bouton volume, interbouton interval, total axon length, maximal horizontal axonal extent, axonal node frequency (/100 µm), and number of primary dendrites were significantly different (Mann-Whitney U Test, * p ≤ 0.05; ** p ≤ 0.01).

However, systematic analysis also revealed a novel group of layer 1 interneurons having large, rosehip-like individual axonal boutons forming very compact, bushy arborizations (rosehip cells, n=10, 13%; Fig. 2A-C). To our knowledge, interneurons having the phenotype of rosehip cells detailed below, have not been identified previously in layer 1 of the cerebral cortex. Somata and dendrites of rosehip cells were confined to layer 1 with only distal dendrites occasionally penetrating layer 2. Proximal dendrites and somata of rosehip cells were decorated with stub-like spines. The axon of rosehip cells usually emerged from the basal part of the soma and gave rise to very compact, dense axonal trees predominantly arborizing in layer 1 with tortuous collaterals having spindle-shaped boutons with diameters not seen in other types of human layer 1 interneurons in our sample. We were unable to find evidence for rosehip cells with recordings performed in layer 2/3. We then systematically increased the number of recorded rosehip cells in our layer 1 database (n=86) and quantitatively compared dendritic and axonal parameters ^24^ of three dimensionally reconstructed rosehip cells (n=6) to neurogliaform cells (n=5; Fig. 2E), the best characterized cell type in the microcircuit of human layer 1 to date ^25,26^. The soma of rosehip cells gave rise to fewer primary dendrites (5.50±1.87) compared to neurogliaform cells (8.6±2.19, n=5, p<0.03, Mann-Whitney (MW) U-test). Total length (11.13±1.99 mm) and maximal horizontal extent of axons (287.75±70.15 µm) of rosehip cells were significantly smaller than those of neurogliaform cells (24.74±8.90 mm, 648.68±202.60 µm, respectively; p<0.004 for both, MW U-test). We measured axonal bouton densities of rosehip (n=6) and neurogliaform (n=4) cells in 10 µm thick spherical shells of increasing diameter by Sholl analysis corrected with the portion of shells outside the brain slice. The bouton density of rosehip and neurogliaform cells monotonously decreased with increasing distances from the soma, however, bouton densities were higher in neurogliaform cells 70-220 µm from the soma (p<0.05 for 70 µm, p<0.01 for 80-220 µm, MW U-test). Rosehip cells had longer interbouton intervals compared to neurogliaform cells (3.97±0.49 and 3.10±0.32 µm, respectively, p<0.038, MW U-test) measured as linear distances between neighboring boutons.

Rosehip cell axons branched more frequently, with rosehip cells and neurogliaform cells having 1.52±0.45 and 0.61±0.21 nodes along 100 µm length of their axons (p<0.004, MW U-test). The axon tortuosity (measured as the average ratio of the actual axonal path and linear distance between neighboring nodes) of rosehip cells (1.42±0.05) was similar to that of neurogliaform cells (1.54±0.15). Measurements based on serial section electron microscopy and three-dimensional reconstructions revealed that the volume of axon terminals of rosehip cells (0.37±0.18 µm^3^, n=31) was approximately four times larger (p<0.001; MW U-test) compared to that of neurogliaform cells (0.08±0.06 µm^3^, n=24, Fig. 2E, 5I). The size of active zones in rosehip axon terminals (0.11±0.03 µm^2^, n=11) was not correlated to bouton volumes (rho=0.34, p=0.29, Spearman correlation).

To understand the molecular identity of rosehip cells and link them to the transcriptomic clusters, we performed immunohistochemistry (IHC) on whole cell recorded and anatomically recovered cells for well characterized antibody markers of GABAergic cell types. This revealed that rosehip cells were immunopositive for CCK (n=6) but negative for CB1 cannabinoid receptor (CNR1, n=5), SST (n=3) and calretinin (CALB2; n=2; Fig. 3A). Furthermore, rosehip cells were immunopositive for gamma-aminobutyric acid (GABA; n=2), and for chicken ovalbumin upstream promoter transcription factor II (NR2F2; n=2), negative for the widely used interneuron marker molecules parvalbumin (n=3), neuronal nitric oxide synthase (n=4), neuropeptide Y (n=2), calbindin (n=2), and choline acetyltransferase (n=3; Suppl. Fig. 2A).

**Figure 3.**
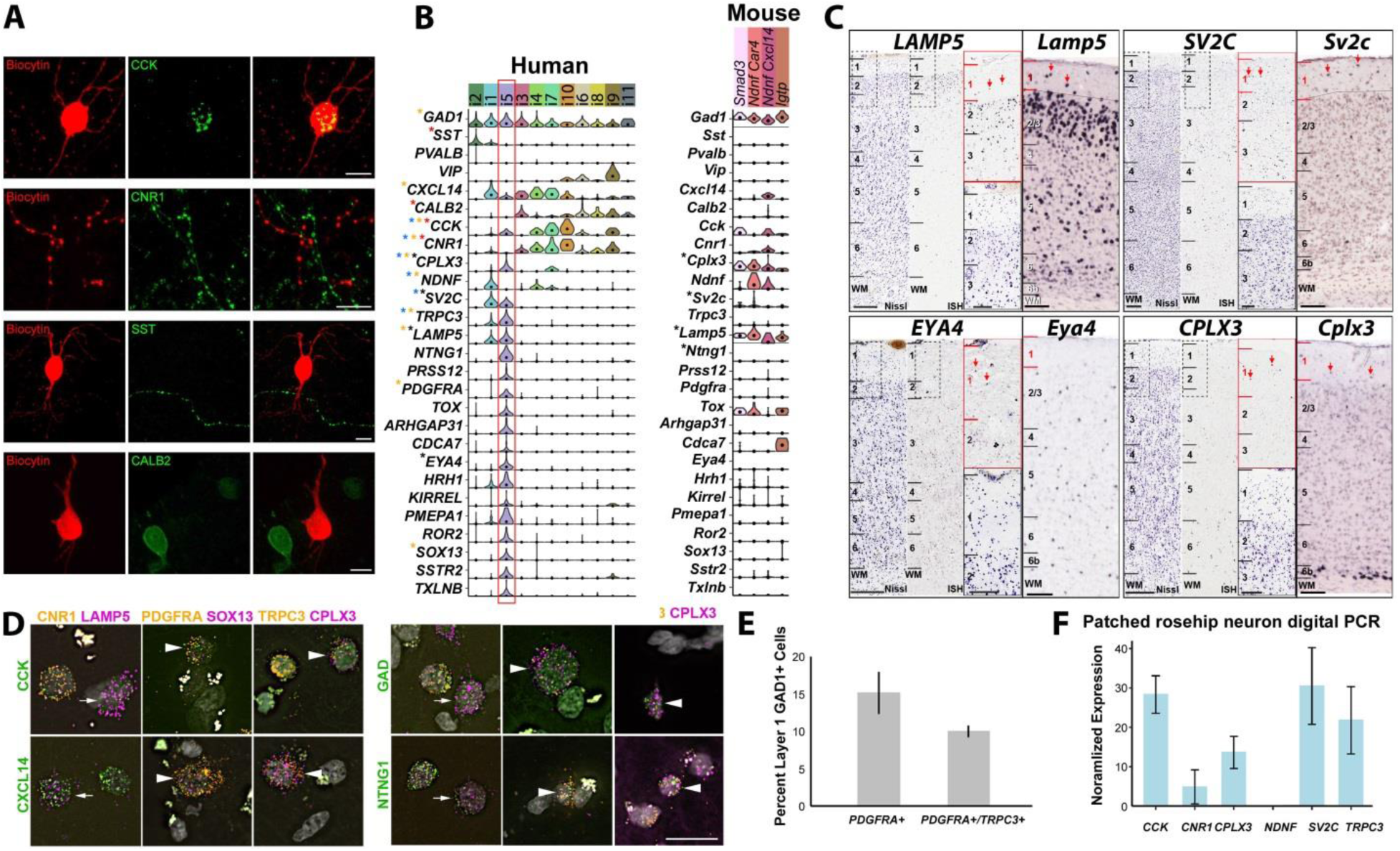
Molecular phenotype of rosehip cells in layer 1 of the human cerebral cortex. **A**, Whole cell recorded and biocytin (red) filled rosehip cells shows CCK (green; n=6) immunopositivity. All biocytin (red) labeled rosehip cells tested for CB1 cannabinoid receptors (CNR1; n=5), somatostatin (SST; n=3), and calretinin (CALB2; n=2) were immuno-negative in spite of having labeled cells in the vicinity. Scale bars, 10 µm. **B**, Violin plots of gene expression for broad cell type and putative rosehip specific markers. Expression validated for select genes by immunohistochemistry (red stars), colorimetric ISH (black), multiplex FISH (orange), and single cell digital PCR (blue) in morphologically identified rosehip cells. **C**, ISH of select marker genes in human temporal cortex (left) and mouse cortex (right). Red arrows highlight cells expressing genes in layer 1. Scale bars=250 µm (low mag), 100 um (high mag). **D**, Multiplex FISH validation of rosehip marker co-expression. Arrowheads and arrows show examples of rosehip cells that are triple- and double-positive (i.e. *CNR1*-), respectively, for marker genes based on RNA-Seq expression data. Scale bar=25 µm. **E**, Rosehip cells comprise 10-15% of layer 1 interneurons based on multiplex FISH quantification of 408 *GAD1*+ cells in 2 subjects. 15% (+/-3) of *GAD1*+ cells express the rosehip specific marker *PDGFRA*, although a small fraction of these cells may be oligodendrocyte precursor cells (see SI Figure 2B). 10% (+/-1) of *GAD1*+ cells express *PDGFRA* and a second rosehip marker *TRPC3*, although some rosehip cells may lack TRPC3 expression based on RNA-seq. Error bars represent standard deviation. **F**, Expression of rosehip cluster markers in cytoplasm of whole cell recorded rosehip cells. Quantified by single cell digital PCR and reported as a percentage of housekeeping gene (*TBP*) expression in 3-4 cells per gene. Note that *NDNF* expression was not detected in any of 3 cells.

Remarkably, this immunohistochemical profile aligned closely with a single transcriptomic cell type, i5, which was similarly GAD1/CCK-positive but CNR1/SST/CALB2/PVALB-negative (Fig. 3B, red box). This putative rosehip transcriptomic type, one of the most distinctive layer 1 GABAergic transcriptomic types, expresses many other genes either highly specific or coexpressed in only one other layer 1 cell type. Intriguingly, given the rosehip synaptic phenotype, these markers include many genes with known associations to axon growth and synaptic structure and function, including synaptic vesicle glycoprotein 2c (*SV2C*), *LAMP5*, transient receptor potential cation channel subfamily C member 3 (*TRPC3*), complexin 3 (*CPLX3*), neurotrypsin (*PRSS12*), netrin G1 (*NTNG1*), histamine receptor H1 (*HRH1*), receptor tyrosine kinase like orphan receptor 2 (*ROR2*), somatostatin receptor 2 (*SSTR2*), and taxilin beta (*TXLNB*).

Since the rosehip synaptic anatomy phenotype has not been described in rodents, we asked whether a transcriptomic signature similar to the rosehip transcriptomic type had been observed in a recently published large-scale analysis of mouse primary visual cortex using single cell RNA-seq analysis ^4^. We did not observe a transcriptional signature even closely resembling the marker gene profile of human rosehip cluster in mouse, either for the PVALB/SST/VIP-negative types (Fig. 3B, right panel) or the complete set of GABAergic types described in that study (Suppl. Fig. 2B). While there were two CCK+/CNR1-clusters in mouse cortex, they did not consistently coexpress the great majority of selective genes seen in the human rosehip cluster. Importantly, it is the unique combinatorial expression of many marker genes that defines rosehip cells. For example, expression of LAMP5, SV2C, EYA4 and CPLX3 is seen by ISH in human layer 1 (Fig. 3C); similarly, as predicted by transcriptomics three of these four genes are also expressed in mouse layer 1 whereas cells expressing *Eya4* are extremely rare. Many other rosehip-selective genes had no evidence of expression in layer 1 interneurons in mouse based on single cell transcriptomics (Fig. 3B, Suppl. Fig. 2B).

To demonstrate that layer 1 neurons with combinatorial expression patterns predicted by transcriptomics could be found in human layer 1, and to quantify their proportions, we systematically performed triple fluorescent ISH using discriminating positive and negative gene markers. For all combinations tested we observed cells with the predicted profiles. For example, we observed CCK+/CNR1-/LAMP5+, CCK+/PDGFRA+/SOX13+, and CCK+/TRPC3+/CPLX3+ cells, as well as cells where CCK was swapped with other positive rosehip markers (Fig. 3D; additional gene combinations shown in Suppl. Fig 2C). Quantification of cell proportions using marker expression is complicated by two factors; first, markers for one cell type are often expressed in others, and second, individual markers are often not expressed in every cell in a cluster. We used the combination of *GAD1*, *PDGFRA* and *TRPC3* to quantify the proportion of rosehip cells among layer 1 GABAergic neurons (Fig. 3E). *PDGFRA* is known to be expressed in OPCs at extremely high levels as well (which is why it appears to only be expressed in OPCs in Figure 1 but appears high in rosehip cells in Figure 2 once levels are not normalized across all cell types including OPCs). *PDGFRA*+ cells represent ~15% of *GAD1*+ cells, therefore an upper bound. *TRPC3* is not expressed in all cells in the rosehip cluster on the other hand. The proportion of GAD1+ cells that are PDGFRA+/TRPC3+ was ~10%, therefore a lower bound. The triple positive cells for this combination were sparsely distributed across layer 1, although not restricted to this layer (Suppl. Fig. 2D).

Finally, to more concretely link morphologically and transcriptionally defined rosehip cells, we performed digital PCR for additional marker genes on cellular content extracted from individual rosehip neurons. As predicted by the transcriptome data, rosehip cells were positive for *CCK*, *CPLX3*, *SV2C* and *TRPC3*, and low (*CNR1*) or absent (*NDNF*) for genes not expressed by cells in that cluster (Fig. 3F). Together, these data strongly link the anatomically-defined rosehip phenotype with a highly distinctive transcriptomic cell type signature that is found in human but not in mouse layer 1.

### Intrinsic electrophysiological properties of rosehip cells

Anatomically identified rosehip cells responded to long (800 ms) suprathreshold current injections with stuttering or irregular spiking firing pattern ^2^ when activated from resting membrane potential (-61.34±5.8 mV, Fig. 4A, 5B,C,E,F,K). Analysis of silent and suprathreshold periods during rheobasic firing of rosehip cells indicated that membrane oscillations and firing of rosehip cells were tuned to beta and gamma frequencies (Fig. 4E-G). The power of averaged fast Fourier transforms (FFT) of subthreshold membrane potential oscillations ^27^ was higher between 3.8 and 80 Hz in rosehip cells compared to neurogliaform and other interneurons (Fig. 4F) and intraburst frequency of stuttering firing also peaked in the beta-gamma range (Fig. 4G). The standard deviation of interspike intervals was higher in rosehip cells (87±64 ms, n=55) compared to neurogliaform (41±34 ms, n=16, p<0.001, Wilcoxon-test) or unclassified (47±41 ms, n=36, p<0.001, Wilcoxon-test) interneurons, indicating alternating silent and active periods during rheobasic stimulation. As described previously, ^25^ human interneurons recorded in layer 1 had a characteristic sag when responding to hyperpolarizing current pulses. However, the amplitude of the sag measured in anatomically classified rosehip cells (1.73±0.30, n=55) exceeded that of interneurons morphologically identified as neurogliaform cells (1.19±0.12, n=16, p<0.001, Wilcoxon-test) or unclassified interneurons (1.29±0.28, n=36, p<0.001, Wilcoxon-test). Input resistances of rosehip cells (139.6±54.1 MΩ) were similar to that of neurogliaform cells (160.1±55.9 MΩ) and lower compared to other interneurons (216.3±84.4 MΩ, p<0.001, Wilcoxon-test), however, time constants of rosehip cells (7.3±3.7 ms) were similar to neurogliaform (8.9±2.4 ms, p<0.001) and faster compared to other cells (11.1±12.5 ms, p<0.001). Anatomically identified rosehip cells showed distinct impedance profiles relative to other layer 1 interneurons in response to current injections with an exponential chirp (0.2-200 Hz, Fig. 4 C-D). Impedance at the lowest frequency (Z_0.2Hz_) was similar in layer 1 interneurons (rosehip, 258±81 MΩ, neurogliaform, 279±128 MΩ, unclassified, 261±133 MΩ, Lillefors test followed by one-way ANOVA with Bonferroni correction). Resonance magnitude (Q, see methods) of rosehip cells (1.77±0.34) was significantly higher compared to neurogliaform cells (1.31±0.07; p<0.021, Lillefors test followed by one way ANOVA with Bonferroni correction) and unclassified interneurons (1.37±0.19; p<0.049). In addition, frequencies of maximal impedance (f_max_) in rosehip cells (4.17±1.1 Hz) were significantly higher than in neurogliaform cells (1.98±1.04 Hz; p<0.045) but the difference was not significant compared to unclassified interneurons (2.47±1.47 Hz, p<0.142). We did not find significant differences between neurogliaform cells and unclassified interneurons in impedance parameters. Support vector machine (SVM) based wrapper feature selection of electrophysiological parameters ranked the amplitude of the sag and the standard deviation of interspike intervals as the two best delineators out of n=200 measured electrophysiological parameters for separating anatomically identified rosehip, neurogliaform and unclassified interneurons in layer 1 (Fig. 4B). Indeed, the best hyperplane separating rosehip cells from other interneuron types according to SVM analysis had a false positive rate of 0% for identifying rosehip cells (n=37) in the total population of anatomically recovered layer 1 interneurons (n=107). Thus, we included cells defined by the hyperplanes of SVM analysis referred to as SVM identified rosehip cells in case anatomical recovery was lacking in some experiments as indicated below.

**Figure 4.**
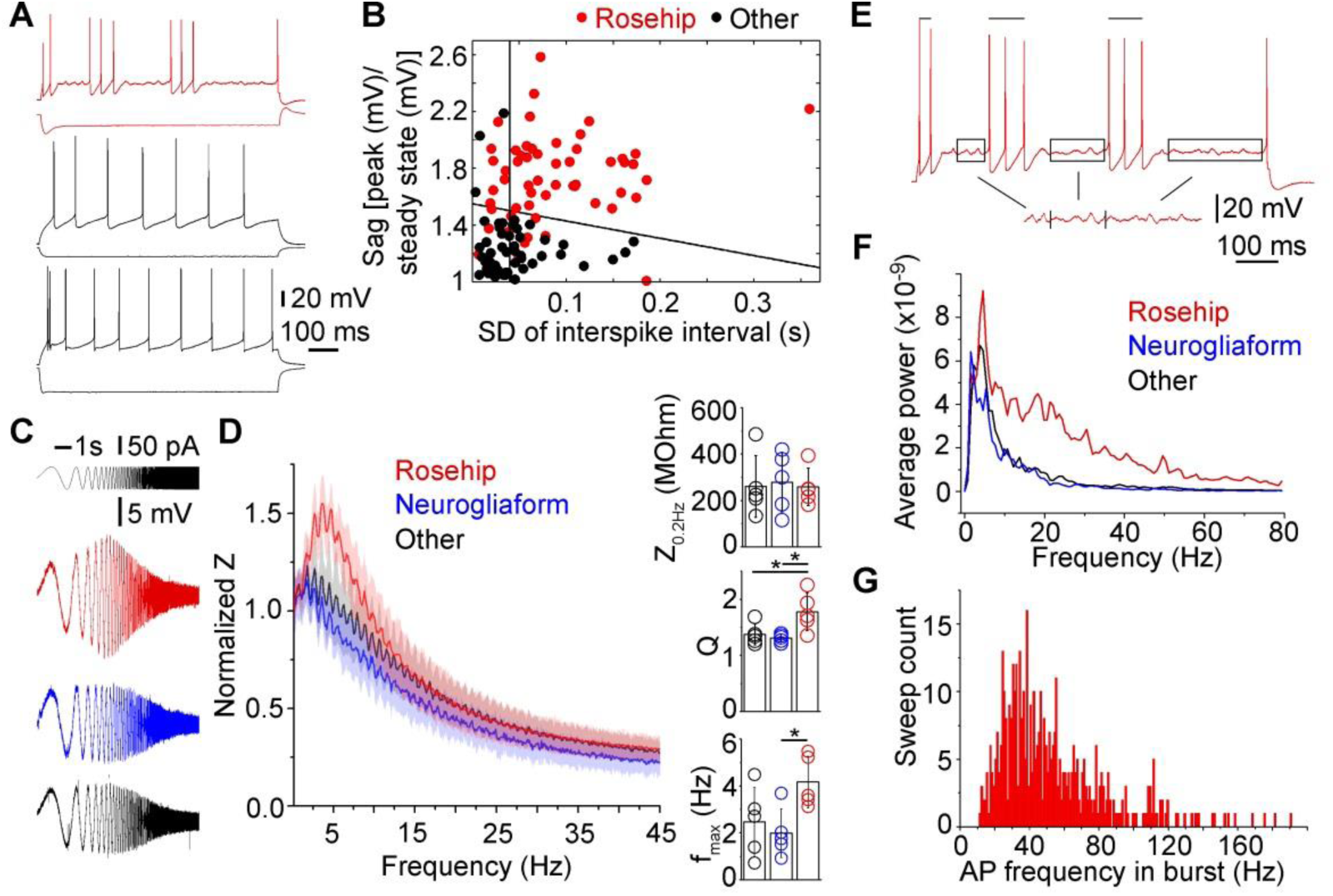
Intrinsic electrophysiological properties of rosehip cells. **A**, Examples of different firing patterns induced by current injections in layer 1 interneurons. Firing pattern of a rosehip cell (top), a neurogliaform cell (middle) and an unidentified layer 1 interneuron (bottom). **B**, Support vector machine (SVM) based wrapper feature selection of electrophysiological parameters for the identification of rosehip cells. Anatomically identified rosehip cells (red dots) and other types of interneurons with known morphology (black dots) are mapped to the distribution of electrophysiological features ranked as the two best delineators by SVM. Black lines show the best hyperplane separating rosehip cells from other interneuron types. **C-D**, Rosehip cells exhibit distinct impedance profile relative to other human interneurons in layer 1. **C**, Responses of anatomically identified rosehip (red), neurogliaform (blue) and other (black) interneurons to current injections with an exponential chirp (0.2-200 Hz, top). Traces were normalized to the amplitude of the rosehip response at 200 Hz. **D**, Left, Normalized impedance (Z) profiles of distinct groups of interneurons. Rosehip cells had higher impedance in the range of 0.9 - 12.4 Hz compared to neurogliaform and other interneurons. Right, Impedances were similar at the lowest frequency (Z_0.2 Hz_, left), but resonance magnitude (Q) calculated as maximal impedance value divided by the impedance at lowest frequency (middle) and frequencies of maximal impedance (f_max_, right) showed significant differences (p<0.05, ANOVA with and Bonferroni post hoc correction). **E**, Automatized selection of recording periods for the assessment of subthreshold membrane potential oscillations (boxed segments) and detection of bursts (bars) for measuring intraburst spiking frequency demonstrated on a rosehip cell response to near rheobasic stimulation showing stuttering firing behavior. **F**, Averaged fast Fourier transforms (FFT) of membrane potential oscillations had higher power between 3.8 and 80 Hz in rosehip cells compared to neurogliaform and other interneurons. **G**, Intraburst frequency of rosehip cells peaked in the gamma range.

**Figure 5.**
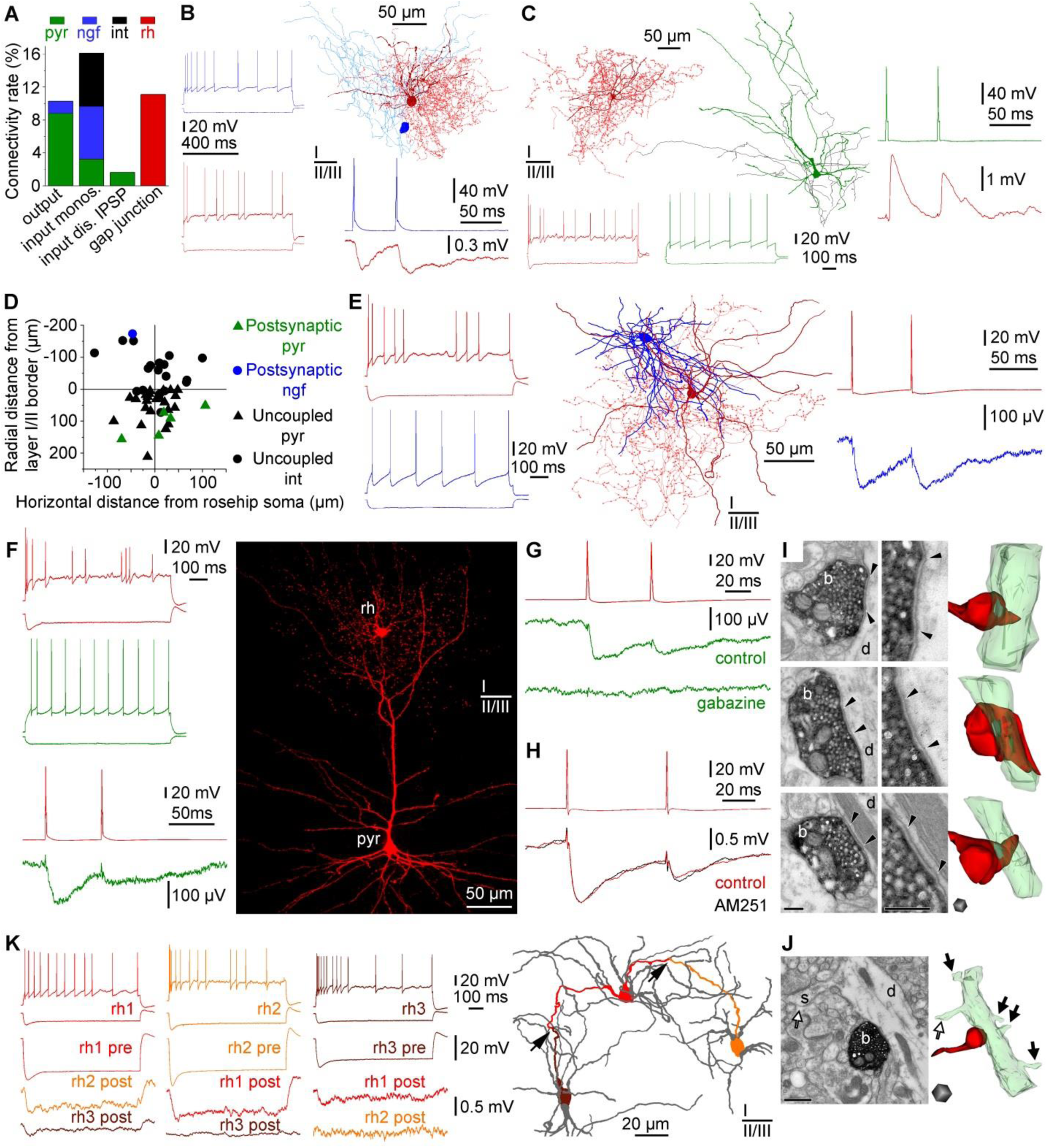
Connections of rosehip cells in the local microcircuit. **A**, Distribution of local connections mapped in layers 1-3 between rosehip cells (rh, red), pyramidal cells (pyr, green), neurogliaform cells (ngf, blue) and other types of layer 1 interneuron (int, black) based on unbiased targeting of postsynaptic cells. Rosehip cells predominantly innervate pyramidal cells, receive monosynaptic EPSPs from layer 2-3 pyramidal cells, monosynaptic IPSPs from neurogliaform and other types of interneurons, however, IPSPs arriving from rosehip cells were not encountered. In addition, rosehip cells are interconnected by homologous electrical synapses (gap junctions). **B**, Example of a neurogliaform cell to rosehip cell connection. Left, Firing patterns of the presynaptic neurogliaform cell (blue) and the postsynaptic rosehip cell (red). Right, Anatomical reconstruction of the recorded neurogliaform cell (soma, dark blue; axon, light blue) and rosehip cell (soma and dendrites, burgundy; axon: red). Action potentials in the neurogliaform cell (blue) elicited slow IPSPs in the rosehip cell (red). **C**, Example of a pyramidal cell to rosehip cell connection. Left, Anatomical reconstruction and firing pattern of the presynaptic pyramidal cell (firing, soma and dendrites, green; axon, black) and the postsynaptic rosehip cell (firing, soma and dendrites, burgundy; axon, red). Right, action potentials in the pyramidal cell (green) elicited EPSPs in the rosehip cell (burgundy). **D**, Spatial distribution of coupled and uncoupled neurons tested as postsynaptic targets of rosehip cells. Note the relative dominance of layer 2-3 pyramidal cells among neurons receiving input from rosehip cells. **E**, The only rosehip cell to neurogliaform cell connection successfully tested for synaptic coupling. Left, Firing patterns of the presynaptic rosehip cell (burgundy) and the postsynaptic neurogliaform cell (blue). Middle, Anatomical reconstruction of the rosehip cell (soma and dendrites, burgundy; axon, red) and the neurogliaform cell (soma and dendrites, blue; axon not shown). Right, Action potentials in the rosehip cell (red) elicited slow IPSPs in the neurogliaform cell (blue). **F**, Example of a rosehip cell to pyramidal cell connection. Left, Firing patterns of the presynaptic rosehip cell (red) and the postsynaptic pyramidal cell (green). Action potentials in the rosehip cell (burgundy) elicited IPSPs in the pyramidal cell (green). Right, Confocal fluorescence image showing the recorded rosehip cell (rh) forming its axonal cloud in the tuft of the apical dendrite of the layer 2-3 pyramidal cell (pyr). **G**, Pharmacological characterization of a rosehip-to-pyramidal cell connection. Presynaptic spikes in the rosehip cell (red) elicited IPSPs in the layer 2-3 pyramidal cell (green) which could be blocked by application of gabazine (10 µM). **H**, Functional test of presynaptic CNR1expression in rosehip cells show the absence of modulation by the CNR1antagonist AM251. Presynaptic spikes in the rosehip cell 1 (red, top) elicited IPSPs in the rosehip cell 2 (red, bottom). Application of AM251 (5 µM) had no effect on IPSPs (black). **I**, Representative electron microscopic images (left) and three-dimensional reconstructions (right) showing axon terminals (b, red) of biocytin filled rosehip cells (n=3) targeting exclusively dendritic shafts (d, green) (100%, n=31). Synaptic clefts are indicated between arrowheads. Scale bars: 200 nm. **J**, Representative electron microscopic image (left) and three-dimensional reconstruction (right) of a biocytin filled rosehip cell bouton (b, red) targeting a pyramidal dendritic shaft (d, green) identified based on emerging dendritic spines (s, arrows). Scale bars: 500 nm. **K**, Rosehip cells form a network of electrical synapses. Top left, firing patterns of three rosehip cells (red, rh1; orange, rh2; burgundy, rh3). Bottom left, Hyperpolarization of rosehip cell rh1 was reciprocally transmitted to rosehip cells rh2 and rh3 confirming electrical coupling. Right, Route of the hyperpolarizing signals through putative dendro-dendritic gap junctions (arrows) between rosehip cells rh1, rh2 and rh3 is shown by corresponding colors in the dendritic network of the three cells (gray).

### Function of rosehip cells in local microcircuits

To assess functional connectivity of rosehip cells in the local microcircuit, we established recordings from rosehip cells and then searched for potential pre- and postsynaptic partners without any cell type preference in an area of the brain slices within a horizontal and vertical radius of ~100 µm and ~200 µm, respectively (Fig. 5A-F). Monosynaptic inputs were tested on anatomically (n=36) and SVM (n=24) identified rosehip cells. Presynaptic layer 1 interneurons were connected to rosehip cells with an overall coupling ratio (CR) of 45%. GABAergic cells evoking IPSPs on rosehip cells included layer 1 neurogliaform cells (n=10, CR 100 %), rosehip cells (n=2, CR 17%) and unclassified interneurons (n=14, CR 40%), however, none of the tested interneurons (n=9) having somata in layer 2 (defined as <70 µm below the layer 1/2 border) were connected to rosehip cells. Fast components of IPSPs arriving to rosehip cells evoked by different presynaptic interneurons had similar amplitudes (0.982±0.705, 0.915±0.594 and 1.504±1.308 mV, respectively) and showed paired pulse depression with paired pulse ratios of (0.42±0.48, 0.27 and 0.71±0.26, respectively). Rosehip cells received local excitatory inputs from layer 2-3 pyramidal cells sporadically (n=7, CR 5%) with monosynaptic EPSP amplitudes of 3.357±1.458 mV and paired pulse ratios of 0.68±0.12. Very large unitary EPSPs, previously described to drive human basket and axo-axonic cells to suprathreshold postsynaptic responses ^10,11,28^, were not encountered on rosehip cells. Thus, acknowledging the fact that some axon collaterals of pyramidal cells were cut during the slicing procedure (Fig. 5C) leading to a potential underrepresentation of pyramidal cell triggered EPSPs, local inputs to rosehip cells appear to be predominantly GABAergic.

In turn, monosynaptic output connections triggered by anatomically (n=41) and SVM (n=13) identified rosehip cells rarely innervated postsynaptic interneurons (overall CR 8%). Even though a neurogliaform cell (n=1, CR 10%), rosehip cells (n=2, CR 17%), unclassified layer 1 interneurons (n=2, CR 5%) and superficial layer 2 pyramidal cells (n=3, CR 4%) were targeted, the output of rosehip cells were predominantly directed towards layer 3 pyramidal cells (n=12, CR 44%) having somata >70 µm below the layer 1/2 border. IPSPs elicited by rosehip cells were mediated by GABAA receptors based on experiments showing blockade of IPSPs by application of the GABAA receptor antagonist gabazine (10 µM, Fig. 5G). Amplitudes of rosehip cell triggered IPSPs arriving to interneurons (0.428±0.370 mV) were larger compared to those targeting layer 3 pyramidal cells (0.087±0.059 mV, p<0.05, MW U-test), in agreement with dendritic filtering of distally elicited IPSPs during signal propagation along the apical dendrite to the somatically placed electrode. The results above indicate that rosehip cells might preferentially target pyramidal cells sending terminal branches of their apical dendrites to layer 1. Indeed, when randomly sampling the output formed by rosehip cells (n=3) using serial electron microscopic sections, we found that axon terminals (n=31) exclusively targeted dendritic shafts (Fig. 5I). Moreover, further ultrastructural analysis of postsynaptic dendrites (n=15) revealed dendritic spines and sparse innervation by symmetrical synapses on the shaft, suggesting that these dendrites belonged to pyramidal cells (n=13, 86%, Fig. 5J). A postsynaptic dendrite (n=1, 7%,) having no spines and receiving asymmetric synapses on the shaft was likely to be formed by an interneuron. Features of the remaining postsynaptic dendrite (n=1, 7%) were insufficient for further identification.

Previous studies on rodent cortical interneurons containing CCK show functional presynaptic expression of the CB1 cannabinoid receptor ^29^, however, application of the CB1 receptor antagonist AM251 was ineffective in modulating rosehip cell evoked IPSPs (n=4, Fig. 5H), supporting our results of single cell digital PCR, IHC and ISH data (Fig. 3). Earlier reports on human microcircuits identified single cell triggered polysynaptic network events ^10,11,28,^. We found that rosehip cells were involved in single cell activated ensembles detected through disynaptic IPSPs triggered by layer 2 (n=1) and layer 3 (n=2) pyramidal cells and polysynaptic EPSPs triggered by an axo-axonic cell, respectively (data not shown). In addition to mono- and polysynaptic chemical synaptic communication, human interneurons are also involved in gap junctional signaling ^25^. Rosehip cells also formed homologous electrical synapses (n=5) between each other (Fig. 5K) and established heterologous electrical synapses (n=1) with an unclassified layer 1 interneuron.

Preferential placement of output synapses on distal dendritic shafts of pyramidal cells reaching layer 1 suggest that rosehip cells might specialize in the control of dendritic signal processing. In dual recordings of synaptically connected rosehip cells to pyramidal cell pairs (n=4), we loaded rosehip cells with Alexa Fluor 594 to label presynaptic axons and filled the postsynaptic pyramidal cells with Oregon Green BAPTA 1 in order to structurally map the course of dendrites and to measure dendritic Ca^2+^ dynamics (Fig. 6). The amplitude of the IPSPs triggered by the first action potential of rosehip cells and evoked on distal dendrites of the postsynaptic pyramidal could be detected at the soma (46.3±27.2 µV) confirming synaptic coupling (Fig. 6A,B). To our knowledge, backpropagation of action potentials to dendrites ^30^ of human neurons has not been addressed in previous studies, thus we tested dendritic Ca^2+^ responses following somatically elicited burst firing (100 ms current injections, 4 spikes/burst) in layer 3 pyramidal cells. Changes in ΔF/F (14.9±5.6%) in distal branches of the apical dendrites in layer 1 were consistently detected at multiple (14±6) locations on the postsynaptic neurons confirming action potential backpropagation into distal apical dendritic branches of human pyramidal cells (Fig. 6C). We chose regions of interest on Oregon Green BAPTA 1-filled branches of the postsynaptic apical dendrites overlapping with the Alexa Fluor 594 labeled axonal arborization of presynaptic rosehip cells and triggered somatically evoked bursts in the pyramidal cells alone for control and together with bursts in the rosehip cell in an alternating fashion (Fig. 6B-F). Inputs from rosehip cells simultaneous with backpropagating action potentials were effective in suppressing the amplitude of Ca^2+^ signals relative to control (n=4, 13.4±3.8% vs. 18.7±3.4% ΔF/F, p<0.02, Wilcoxon-test, Fig. 6C) in one or two locations heuristically line scanned on dendrites of postsynaptic cells (Fig. 6G). The anatomical arrangement of presynaptic axons and imaged segments of postsynaptic dendrites was recovered in n=3 pairs. Rosehip inputs simultaneous with backpropagating pyramidal action potentials were effective in suppressing Ca^2+^ signals only at sites that were neighboring (10±3 µm) to the putative synapses between the two cells. No effect of rosehip cells was detected at dendritic sites one step further in distance (23±12 µm, Fig. 6D-F). This suggests that rosehip cells specialize in providing tightly compartmentalized control of dendritic Ca^2+^ electrogenesis of human pyramidal cells enforcing inhibitory microdomains in dendritic computation.

**Figure 6.**
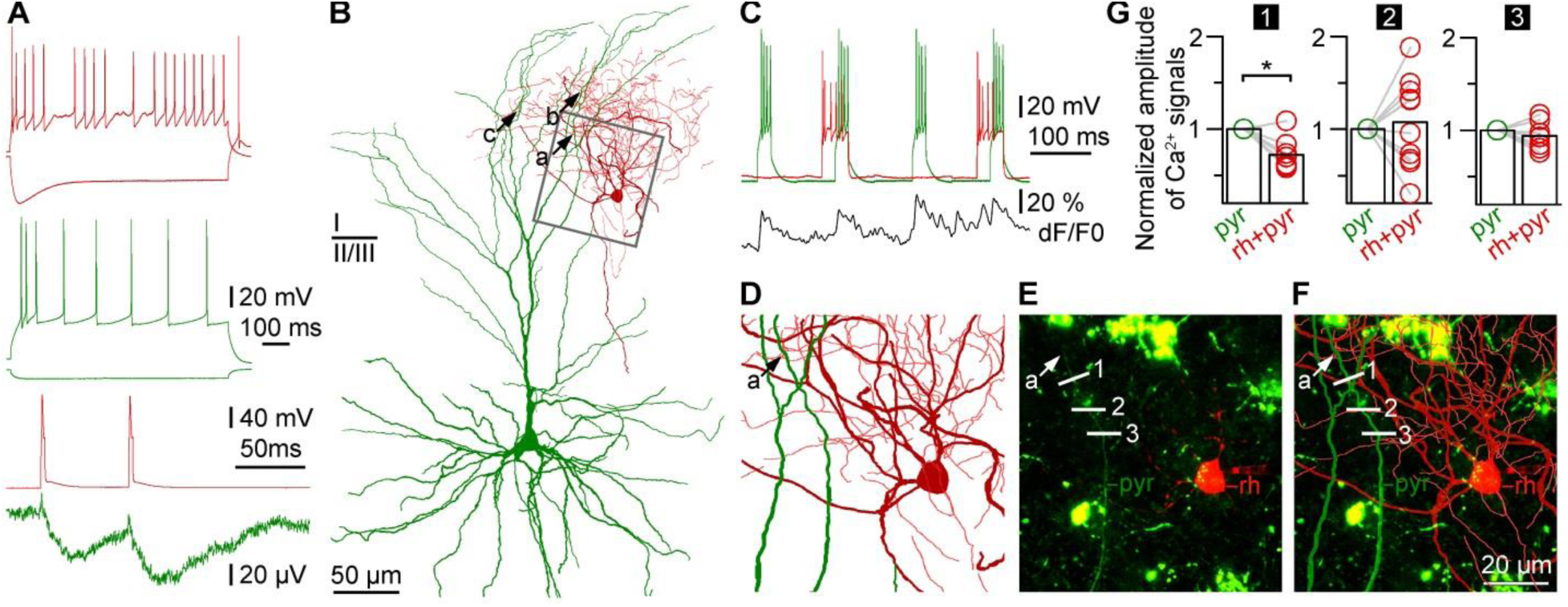
Human rosehip interneurons perform segment specific regulation of action potential backpropagation to apical dendritic tufts of pyramidal cells. **A**, Top, Firing patterns of a presynaptic rosehip cell (burgundy) and a postsynaptic pyramidal cell (green). Bottom, Action potentials in the rosehip cell (burgundy) elicited IPSPs in the pyramidal cell (green). **B**, Anatomical reconstruction of the rosehip cell (soma and dendrites: burgundy; axon, red) and the layer 2-3 pyramidal cell (soma and dendrites, green; axon not shown). Presynaptic axonal boutons of the rosehip cell formed close appositions (a, b, and c) with three separate branches on the tuft of the pyramidal apical dendrite. **C**, Repetitive burst firing was triggered to initiate backpropagating Ca^2+^ signals in the pyramidal cell (green) while the output of the rosehip cell (red) was switched on and off timed prior and during every second pyramidal burst. Simultaneously, Ca^2+^ dynamics of the pyramidal apical dendritic tuft was measured at several locations and signals detected at location no.1 shown on panels E and F are shown in black. **D**, The area boxed in panel B shows the dendritic branch of the apical tuft of the pyramidal cell (green) with a putative synaptic contact (a) arriving from the rosehip cell. **E**, Confocal Z-stack image of the same area shown on panel D taken during paired whole cell recordings. The soma of the rosehip cell (rh, red), the dendrite of the pyramidal cell (pyr, green), the putative synaptic contact (a) arriving from the rosehip cell to the pyramidal cell and sites of line scans performed across the dendrite (1, 2 and 3) are indicated. Cytoplasmic lipofuscin autofluorescence characteristic to human tissue is seen as green patches. **F**, Superimposition of the anatomical reconstruction of panel D and the confocal image of panel E. **G**, Normalized amplitudes of Ca^2+^ signals during pyramidal cell firing with and without coactivation of the rosehip cell detected at the three sites of line scans (1, 2 and 3) on the pyramidal dendrite. Rosehip input simultaneous with the backpropagating pyramidal action potentials was significant (p=0.02) in suppressing Ca^2+^ signals only at site 1 which was closest (8 µm) to the putative synapse between the two cells, no effect (p=1 and p=0.27, respectively) of the rosehip cell was detected at sites 2 and 3 located at distances of 21 and 28 µm, respectively from the putative synaptic contact.

## Discussion

Understanding the cellular makeup of the cortex and its conservation across species represent twin challenges difficult to address in human tissue. Historically, forming a representative overview of cell type diversity in a particular brain region has been achieved based on molecular marker expression cross referenced to axonal and dendritic morphology ^3,4,20,31–33^. Many conserved patterns of molecular expression and morpho-physiological features for a given cell type or class have been reported ^2,34^, but species variation has also been documented ^26,35–39^ and cell types potentially characteristic to a number of species have also been described ^40–43^. Importantly, recent studies have overcome some of the difficulties associated with the scarcity of human tissue of sufficient quality ^8–13,21,25,26,28,35,39,44–50^ paving the way for an in depth understanding of human circuits. Here we demonstrate the strength of a modern version of this approach that can be applied to human postmortem and neurosurgical tissues. Single nucleus transcriptomics provides the scale for an unbiased survey of molecular expression, while in vitro human tissue physiology characterizes the functional properties of those types. Together these approaches provide convergent evidence for a robust description of cell type identity as well as multiple lines of evidence for species conservation or specialization.

The targeted application of single nucleus sequencing reported here has demonstrated a significantly higher degree of GABAergic neuron complexity in just one layer of the human cerebral cortex (10 types) than what has previously been described in all of the cortex (8 types, ^51^). This difference is likely due to a combination of improved sequencing technique and increased sampling in a targeted anatomical domain enriched in GABAergic neurons. This diversity also appears to be higher than described for layer 1 in mouse ^4^, although that study did not specifically target layer 1 and therefore is likely undersampled compared to the current study. On the other hand, a recent systematic characterization of rat cortex ^3^ described 6 morphological types in layer 1 and 17 morpho-electric types, so our results are roughly consistent with the neuronal diversity described using other methods. The rosehip cell represents a type with highly distinctive transcriptomic signature, a highly distinctive morphological, physiological and connectional phenotype, and a strong correspondence between these properties. In this respect, it appears similar to other highly specialized and distinctive cortical cell types such as chandelier cells ^52,53^. Supporting this correspondence between transcriptional and anatomical phenotypes, many of the most selective genes associated with rosehip cells relate to synaptic structure and function. To our knowledge a similar anatomical cell type has not been described in rodent. While we cannot prove the negative, given the enormous focus on cellular studies of rodent cortex such cells would have to be either extremely rare or experimentally difficult to study to have escaped detection to date. Similarly, the rosehip molecular signature appears highly distinctive from any published data from rodent. Although the transcriptomic comparison is between human temporal cortex and mouse visual cortex, regional differences seem unlikely to account for this difference as we found the anatomically defined rosehip type in multiple regions of human cortex. A complete comparison of all cortical cell types and assessment of relative similarities between cell types should be possible in the future as more comprehensive transcriptome data become available and linked to other cellular phenotypes in multiple species.

It is widely accepted ^29^ that CCK-positive cells in the rodent show selectively high expression of cannabinoid receptors and are involved in perisomatic inhibition. The single axonal bouton morphology and/or the compact axonal field of rosehip cells resembles that of cell types described in deeper layers of the cat cortex that innervate relatively proximal dendrites (dendrite targeting and clutch cells) ^54,55^. In contrast, rosehip cells are CCK-positive but cannabinoid receptor-negative, and appear to selectively target distal dendrites of pyramidal neurons. Moreover, when assessing layer 1 canonical inhibitory pathways in rodent with high throughput electrophysiology capable of sampling all cell types in layer 1,Lee at al. ^56^ found two interneuron types and two canonical pathways involving feed forward interneuron-to-interneuron connections. Thus, the monosynaptic pyramidal cell preferring pathway initiated by rosehip cells does not appear to be a viable concept in the rodent layer 1 circuit. Furthermore, focal intralayer inhibition restricted by the compact axonal arbor of rosehip cells to distal dendrites of a column of pyramidal cells is also missing from the rodent; rather, mouse feedforward inhibitory connections are vertically spread to all somatodendritic domains ^56^.

The addition of new human cell types, or specialization of existing types through major modification of cellular features, to cortical networks would be expected to alter circuit function ^3,57,58^, and therefore cannot be studied in rodents. Rosehip cells may be of particular importance in compartmental control of backpropagating action potentials and their pairing with incoming excitatory inputs. The uniquely small membrane capacitance (C_m_) found in human pyramidal cells ^8^ promotes backpropagation of action potentials ^30^ and increases excitability in human dendrites relative to rodent dendrites having larger C_m_. As demonstrated here, action potentials backpropagate to distal dendrites of human pyramidal cells and can be attenuated by rosehip cell activation. Thus, they may represent a mechanism for supplementary inhibitory control required to balance the potentially higher excitability in human dendrites ^8^ and might form the basis of spatially accurate modulation of interactions between long range excitatory inputs arriving to layer 1 and backpropagating action potentials suggested to participate in interhemispheric modulation ^59^. The sharp resonance in the theta-range detected in individual rosehip cells and its potential spread through gap junctions to a rosehip network could phase-selectively interact with long range inputs similarly to mechanisms suggested for example in oscillation dependent memory consolidation ^31,60^.

There is great promise in convergent transcriptomic, anatomical and functional studies in human cortex to establish which features are conserved and divergent among mammals, and these studies are now feasible. The function of neuron types specific to the human circuit could be important in understanding pathological alterations of network functions. For example, several highly selective markers for rosehip cells have been implicated as risk factors for neuropsychiatric disease, including netrin G1 (NTNG1) for Rett syndrome ^61^ and neurotrypsin (PRSS12) for mental retardation ^62^. A better understanding of human cellular and circuit organization may help counteract the current lack of success in translating promising rodent results to effective treatment against human neuropsychiatric disorders ^63,64^.

## Methods

### Human brain specimens

After obtaining permission from decedent next-of-kin, postmortem adult human brain tissue was collected by the San Diego Medical Examiner’s office and provided to the Allen Institute for Brain Science. All tissue collection was performed in accordance with the provisions of the Uniform Anatomical Gift Act described in Health and Safety Code §§ 7150, et seq., and other applicable state and federal laws and regulations. The Western Institutional Review Board reviewed tissue collection processes and determined that they did not constitute human subjects research requiring IRB review. The tissue specimens used in this study were pre-screened for known neuropsychiatric and neuropathological history, and underwent routine serological testing and toxicology screening. Specimens were further screened for RNA quality and had an RNA integrity number (RIN) ≥7. The specimens used in this study were from two individual control Caucasian male donors, aged 50 and 54 years. Postmortem interval was 24 hours for both specimens.

### Tissue processing

Whole postmortem brain specimens were bisected through the midline and individual hemispheres were embedded in alginate for slabbing. Coronal brain slabs were cut at 0.5-1cm intervals through each hemisphere and the slabs were frozen in a bath of dry ice and isopentane and stored at −80°C. Middle temporal gyrus (MTG) was identified on slabs of interest and removed for further sectioning. MTG tissue was then thawed in a buffer containing PBS supplemented with 10mM DL-Dithiothreitol (DTT, Sigma Aldrich), mounted on a vibratome (Leica), and sectioned at 500µm in the coronal plane. Sections were transferred to a fluorescent Nissl staining solution (Neurotrace 500/525, ThermoFisher Scientific) prepared in PBS with 10mM DTT and 0.5% RNasin Plus RNase inhibitor (Promega). After staining for 5 min, sections were visualized on a fluorescence dissecting microscope (Leica) and layer 1 was microdissected using a needle blade micro-knife (Fine Science Tools).

### Nuclei isolation and FACS

Microdissected sections of layer 1 from MTG were transferred into nuclei isolation medium containing 10mM Tris pH 8.0 (Ambion), 250mM sucrose, 25mM KCl (Ambion), 5mM MgCl_2_ (Ambion) 0.1% Triton-X 100 (Sigma Aldrich), 1% RNasin Plus, 1X protease inhibitor (Promega), and 0.1mM DTT and placed into a 1ml dounce homogenizer (Wheaton). Tissue was homogenized to liberate nuclei using 10 strokes of the loose dounce pestle followed by 10 strokes of the tight pestle. Homogenate was strained through a 30µm cell strainer (Miltenyi Biotech) and centrifuged at 900xg for 10 min to pellet nuclei. Nuclei were then resuspended in staining buffer containing 1X PBS (Ambion), 0.8% nuclease-free BSA (Omni-Pur, EMD Millipore), and 0.5% RNasin Plus. Mouse monoclonal anti-NeuN antibody (EMD Millipore) was applied to nuclei preparations at a concentration of 1:1000 and samples were incubated for 30 min at 4°C. Control samples were incubated with mouse IgG1,k isotype control (BD Pharmingen). Samples were then centrifuged for 5 min at 500xg to pellet nuclei and pellets were resuspended in staining buffer as described above. Nuclei samples were incubated with secondary antibody (goat anti-mouse IgG, Alexa Fluor 594, ThermoFisher Scientific) for 30 min at 4°C, centrifuged for 5 min at 500xg, and resuspended in staining buffer. DAPI (4′, 6-diamidino-2-phenylindole, ThermoFisher Scientific) was applied to nuclei samples at a concentration of 1µg/ml.

Single nucleus sorting was carried out on a BD FACS Aria Fusion instrument (BD Biosciences) using a 130µm nozzle. Nuclei were first gated on DAPI and then passed through doublet discrimination gates prior to being gated on NeuN (Alexa Fluor 594) signal. Approximately 10% of nuclei were intentionally sorted as NeuN-negative to allow for the collection of non-neuronal nuclei. Single nuclei were sorted into 96-well PCR plates (ThermoFisher Scientific) containing 2µl of lysis buffer (0.2% Triton-X 100, 0.2% NP-40 (Sigma Aldrich), 1U/µl RNaseOut (ThermoFisher Scientific), PCR-grade water (Ambion) and ERCC spike-in synthetic RNAs [Ambion]). 96-well plates containing sorted nuclei were then snap frozen and stored at −80°C. Positive controls (10 nuclei pools and/or 10 pg and 1 pg total RNA) were included on every 96-well plate of sorted nuclei.

### cDNA and sequencing library preparation

cDNA libraries from single nuclei were prepared using Smart-seq2 ^65^ with minor modifications. Briefly, Protoscript II (New England Biolabs) was used for reverse transcription, the final dilution of ERCCs in the reverse transcription reaction was 1:55 million, and the template switching oligonucleotide was 5’-biotinylated. Additionally, the number of PCR cycles used for cDNA amplification was increased to 21 to compensate for lower RNA content in single nuclei. cDNA yield was quantified using PicoGreen (ThermoFisher Scientific) and a subset of single nuclei libraries were screened for quality on a Bioanalyzer (High Sensitivity DNA Chip, Agilent Technologies). cDNA library quality was further assessed using qPCR for a housekeeping gene (ACTB) and an ERCC spike-in control RNA (ERCC-00009) ^66^.

Sequencing libraries were prepared using Nextera^®^ XT (Illumina) with minor modifications. Briefly, the input amount of cDNA was 250pg per reaction, reactions were carried out a 1/4X the volume recommended by the manufacturer, and the tagmentation step was extended to 10 min. Sequencing library concentration was determined using PicoGreen and 53-57 samples were pooled per sequencing lane. Pooled libraries were purified using Ampure XP beads and eluted to a concentration of 5nM. Following purification, the pooled library size using a Bioanalyzer and Kapa Library QC was used to determine nM concentrations. Final library pools were then diluted to 3nM final concentration. Pooled samples were sequenced on a HiSeq^®^ 4000 instrument (Illumina) using 150 base paired end reads at a mean untrimmed read depth of ~19 million reads/sample and a mean trimmed read depth of ~16 million reads/sample.

### RNA-Seq processing

The RNASeq data obtained from single nuclei is processed and analyzed according to the procedure described in detail previously ^66^. Briefly, following the demultiplexing of the barcoded reads generated on the Illumina HiSeq platform, the amplification (cDNA & PCR) and sequencing primers (Illumina) and the low quality bases were removed using the Trimmomatic software package ^67^. The trimmed reads were mapped to the human reference genome version, GRCh38 (Ensembl) guided by the version 21 annotations obtained from the GENCODE repository. RSEM ^68^ and TOPHAT-CUFFLINKS ^69^ were used to quantify transcript expression at the transcriptome (exon) and the whole genome (exon plus intron) level, respectively. The fastQC (http://www.bioinformatics.babraham.ac.uk/projects/fastqc/), FASTX (http://hannonlab.cshl.edu/fastx_toolkit/download.html), RSeQC ^70^, RNA-seq-QC ^71^ programs were used to generate various sequence and alignment quality metrics used for classification of the sample quality. A novel pipeline (“SCavenger”, unpublished) was created to automate execution across statistical analysis tools, integrate pre-formatted laboratory and clustering metrics, and calculate new statistics specific to biases identified in the single nuclei lab and sequence preparation protocol. The normalized expression counts (FPKM/TPM) generated at both gene and isoform level by RSEM and TOPHAT-CUFFLINKS analyses and the raw counts generated from the RSEM/TOPHAT alignment (BAM) files by the HTSeq-count program ^72^ were used for differential expression analysis.

### RNA-Seq quality control

To remove data from low quality nuclei samples prior to downstream analysis, we implemented a Random Forest machine learning classification approach as described in detail in Aevermann et al. ^73^. The overall workflow for sample quality classification and filtering was to i) establish a training set using a representative subset of samples, ii) collect a series of 108 quality control metrics (e.g. percent unique reads, percent reads surviving trimming, transcript isoform counts) spanning both the laboratory and data analysis workflows as model features, iii) use these training data and quality control metrics to build a classification model using the Random Forest method, and iv) apply the model to the entire data set for quality classification and data filtering.

A training set of 196 samples, including 169 single nuclei samples, was selected and a set of high confidence pass/fail calls for individual samples determined based on the qualitative assessment of data produced by fastQC, which includes quality Phred scores, GC content, Kmer distributions, and sequence over-representation information. Pass samples (152 samples, including single nuclei and purified bulk RNA positive controls) were identified as having high average quality per read across the entire length of the sequenced fragment and a unimodal average GC content around 40%, reflecting the GC content of the expressed human transcriptome. In contrast, two types of Fail samples were identified. One type of Fail samples (29 samples) exhibited a significant number of reads with low mean Phred quality, and average Phred quality scores that fall off down the length of the sequence read. A second type of Fail samples (15 samples) showed a second peak in the GC content distribution with a mean around 48% GC; this peak appears to be generated from ERCC reads, which are derived from bacterial genome sequences.

The quality control metrics for these training data were then used as features to construct a Random Forest model to distinguish these three quality classes (Pass, Fail-Phred, and Fail-ERCC) comprised of one hundred thousand decision trees generated by standard bagging methods as implemented in KNIME v3.1.2. Using this Random Forest classification model, all 196 samples in the training set were classified correctly with high confidence scores. To test the classification accuracy of the resulting random forest model, we used an independent test set of 185 single nuclei samples classified using the same fastQC evaluation criteria applied to the training data, with 135 determined to be Pass samples, 29 determined to be Fails and 21 determined to be Marginals. Application of the random forest model to these test Pass and Fail samples resulted in only 8 misclassifications (4.9%), for a classification accuracy of 95%. The Random Forest model was then applied to the remaining data and final classification determined. A Pass confidence cutoff of 0.6 or greater was used to select single nuclei data for downstream analysis. Using this Random Forest model applied to the entire Layer 1 dataset including contaminating layer 2 excitatory and inhibitory nuclei, 79% of 1154 single nuclei samples passed quality control. For these Pass samples, the average number of reads after trimming was 16,383,881 (±19,810,661), the number of ERCC transcripts detected was 43.76 (± 3.77), and the number of genes detected at a level of >1 FPKM was 6337 (± 1659), giving an average coverage of 879 reads per human gene detected.

### Gene expression calculation

For each nucleus, expression levels were estimated based on the scaled coverage across each gene. Specifically, bam files were read into *R* ^74^ using the “readGAlignmentPairs” function in the “GenomicAlignments” library, and genomic coverage was calculated using the “coverage” function in “GenomicRanges”^75^. All genes in GENCODE human genome GRCh38, version 21 (Ensembl 77; 09-29-2014) were included, with gene bounds defined as the start and end locations of each unique gene specified in the gtf file (<u>https://www.gencodegenes.org/releases/21.html</u>). Total counts for each gene (including reads from both introns and exons) were estimated by dividing total coverage by twice the read length (150bp, paired end). Expression levels were normalized across nuclei by calculating counts per million (CPM) using the “cpm” function in “edgeR” ^76^.

### Clustering nuclei

Nuclei that passed quality control were grouped into transcriptomic cell types based on an iterative clustering procedure. For each gene, log2(CPM + 1) expression was centered and scaled across nuclei. Gene expression dropout was more likely to occur in nuclei with lower quality cDNA libraries and for genes with lower average expression in nuclei isolated from the same cell type. Expression noise models were estimated for each nucleus based on the 8 most similar nuclei using the “knn.error.models” function of the “scde” R package as described in ^77^. These noise models were used to select significantly variable genes (adjusted variance > 1.25) and to estimate a zero-weight matrix that represented the likelihood of dropouts based on average gene expression levels. Dimensionality reduction was performed with principal components analysis (PCA) on variable genes, and the covariance matrix was adjusted by the zero-weight matrix to account for gene dropouts. Principal components (PCs) were retained for which more variance was explained than the broken stick null distribution or PCs based on permuted data. If more than 2 PCs were retained, then dimensionality was further reduced to 2-dimensions using t-distributed stochastic neighbor embedding (tSNE)^78^ with a perplexity parameter of 80.

After dimensionality reduction, nuclei were clustered using a conservative procedure that attempted to split them into the fewest number of clusters possible. Nearest-neighbor distances between all nuclei were calculated and sorted, and segmented linear regression (R package “segmented”) was applied to estimate the distribution breakpoint to help define the distance scale for density clustering. Next, density clustering (R package “dbscan” ^79^) was applied to nuclei, and the number of clusters calculated for a range of 10 nearest-neighbor distances (parameter epsilon), starting from the maximum distance between nuclei to the distance breakpoint identified in the last step. If only one cluster was found using all values of epsilon, then the above procedure was repeated using a perplexity parameter of 50, 30, and 20 for tSNE, and stopping when more than one cluster was detected. Finally, if no cluster splitting was possible using tSNE, then a final density clustering was applied to the first two significant PCs. If more than one cluster was identified, then the statistical significance of each cluster pair was evaluated with the R package “sigclust” ^80^, which compares the distribution of nuclei to the null hypothesis that nuclei are drawn from a single multivariate Gaussian. Iterative clustering was used to split nuclei into sub-clusters until the occurrence of one of four stop criteria: 1) <6 nuclei in a cluster (because it cannot be split due a minimum cluster size of 3), 2) no significantly variable genes, 3) no significantly variable PCs, 4) no significant sub-clusters.

To assess the robustness of clusters, the iterative clustering procedure described above was repeated 100 times for random subsamples of 80% of nuclei. A co-clustering matrix was generated that represented the proportion of clustering iterations that each pair of nuclei were assigned to the same cluster. Average-linkage hierarchical clustering was applied to this matrix followed by dynamic branch cutting (R package “WGCNA”) with cut height ranging from 0.01 to 0.99 in steps of 0.01. A cut height resulting in 25 clusters was selected to balance cohesion (average within cluster co-clustering) and discreteness (average between cluster co-clustering) across clusters. Finally, gene markers were identified for all cluster pairs, and clusters were merged if they lacked binary markers (gene expressed in >50% nuclei in first cluster and <10% in second cluster) with average CPM > 1 (see also Marker gene selection below).

### Cluster visualization

The relationships between cell type clusters were represented as a constellation diagram where the area of each disc is proportional to the number of nuclei in each cluster and the similarity between clusters is proportional to the width of the lines connecting clusters. Cluster similarity was calculated as the average co-clustering between all nuclei for each pair of clusters. For example, a similarity of 0.1 indicates that 1 out of 10 clustering iterations nuclei from one cluster were assigned to the other cluster. Similarities less than 0.05 were not plotted.

Next, clusters were arranged by transcriptomic similarity based on hierarchical clustering. First, the average expression level of each gene was calculated for each cluster. Genes were then sorted based on variance and the top 2000 genes were used to calculate a correlation-based distance matrix, Dxy=1-(cor(x,y))/2, between each cluster average. A cluster tree was generated by performing hierarchical clustering on this distance matrix (using “hclust” with default parameters), and then reordered to show inhibitory clusters first, followed by excitatory clusters and glia, with larger clusters first, while respecting the tree structure. Note that this measure of cluster similarity is complementary to the co-clustering similarity described above. For example, two clusters with high transcriptomic similarity but a few distinct marker genes may have low co-clustering similarity.

### Marker gene selection

Initial sets of marker genes for each pair of clusters were selected by assessing significance of differential expression using the “limma” ^81^ R package, and then filtering these sets of significant genes to include only those expressed in more than 50% of nuclei in the “on” cluster and fewer than 20% of nuclei in the “off” cluster. Potential marker genes for individual clusters were chosen by ranking the significance of pairwise marker genes, summing the ranks across all possible pairs for a given cluster, and sorting the resulting gene list ascending by summed rank. The final set of marker genes was selected by comparing the gene expression distribution for the top ranked marker genes for each cluster using the visualization described below.

### Gene expression visualization

Gene expression (CPM) was visualized using heat maps and violin plots, which both show genes as rows and nuclei as columns, sorted by cluster. Heat maps display each nucleus as a short vertical bar, color-coded by expression level (blue=low; red=high), and clusters ordered as described above. The distribution of marker gene expression across nuclei in each cluster were represented as violin plots, which are density plots turned 90 degrees and reflected on the Y-axis. Black dots indicate the median gene expression in nuclei of a given cluster; dots above Y=0 indicate that a gene is expressed in more than half of the nuclei in that cluster.

### Colorimetric *in situ* hybridization

*In situ* hybridization data for human temporal cortex and mouse cortex was from the Allen Mouse Brain Atlas ^38^ and a comparable study in human temporal cortex ^82^. All data is publicly accessible through <u>www.brain-map.org</u>. Data was generated using a semiautomated technology platform as described ^38^. with modifications to work with postmortem human tissues as described in ^82^. Digoxigenin-labeled riboprobes were generated for each human gene such that they would have >50% overlap with the orthologous mouse gene in the Allen Mouse Brain Atlas ^38^. Mouse ISH data shown is from the region most closely corresponding to human temporal cortex, corresponding to the medial portion of TeA in Paxinos Atlas ^83^.

### Multiplex fluorescent in situ hybridization

The RNAscope multiplex fluorescent kit was used according to the manufacturer’s instructions for fresh frozen tissue sections (ACD Bio), with the exception that fixation at 4 C with 4% PFA was performed for 60 minutes on 20 µm human brain sections, and the protease treatment step was shortened to 15 min. Probes used to identify specific cell types in layer 1 were designed antisense to the following human genes: CCK (hs-539041), CNR1 (hs-591521), CPLX3 (hs-487681-C3), GAD1 (hs-404031 and hs-404031-C3), LAMP5 (hs487691-C3), SV2C (hs448361-C3), PRSS12 (hs-493931-C3), SOX13 (hs-493941-C3), TRPC3 (hs-427641-C2), NTNG1 (hs-446101), CXCL14 (hs-425291), PDGFRA (hs-604481-C2), SOX9 (hs-404221-C2). Positive controls (POLR2A, UBC and PPIB) were used on each tissue sample to ensure RNA quality (ACD Bio, 320861). Following hybridization and amplification, FISH sections were imaged using a 40X oil immersion lens on a Nikon TiE fluorescent microscope. RNA spots in each channel were quantified manually using the ImageJ cell counting plug-in. To count the percentage of Rosehip cells in layer 1, GAD1+ cells were first identified, followed by the PDGFRA+ cells within that population, followed by the TRPC3+ cells in that population. These counts were used to calculate the percentage of the GAD1+ cells expressing PDGFRA and TRPC3. A total of 408 GAD1+ cells were identified from two individuals for this quantification.

### Electrophysiological recordings

All procedures were performed according to the Declaration of Helsinki with the approval of the University of Szeged Ethical Committee. We used neocortical tissue surgically removed from patients (n=32, n=18 female and n=14 male, aged 47±16 years) in a course of five years as part of the treatment protocol for aneurysms (n=7) and brain tumors (n=25). Anesthesia was induced with intravenous midazolam and fentanyl (0.03 mg/kg, 1– 2 lg/kg, respectively). A bolus dose of propofol (1–2 mg/kg) was administered intravenously. To facilitate endotracheal intubation, the patient received 0.5 mg/kg rocuronium. After 120 seconds, the trachea was intubated and the patient was ventilated with a mixture of O_2_-N_2_O at a ratio of 1:2. Anesthesia was maintained with sevoflurane at monitored anesthesia care (MAC) volume of 1.2–1.5. Tissue blocks were removed from prefrontal (n=16), temporal (n=6) and parietal (n=10) areas. Blocks of tissue were immersed in ice-cold solution containing (in mM) 130 NaCl, 3.5 KCl, 1 NaH_2_PO_4_, 24 NaHCO_3_, 1 CaCl_2_, 3 MgSO_4_, 10 d(+)-glucose, saturated with 95% O_2_ and 5% CO_2_ in the operating theatre. Slices were cut perpendicular to cortical layers at a thickness of 350 µm with a vibrating blade microtome (Microm HM 650 V) and were incubated at room temperature for 1 h in the same solution. The solution used during recordings differed only in that it contained 2 mM CaCl_2_ and 1.5 mM MgSO_4_. Somatic whole-cell recordings were obtained at approximately 36 ºC from up to four concomitantly recorded cells visualized by infrared differential interference contrast videomicroscopy at depths 60–130 µm from the surface of the slice. Signals were filtered at 8 kHz, digitized at 16 kHz, and acquired with Patchmaster software. Micropipettes (5–7 MΩ) were filled with a low [Cl]i solution containing (in mM) 126 K-gluconate, 4 KCl, 4 ATP-Mg, 0.3 GTP-NA2, 10 HEPES, 10 phosphocreatine, and 8 biocytin (pH 7.20; 300 mOsm). Presynaptic cells were stimulated with brief (2–10 ms) suprathreshold pulses delivered at >7-s intervals, to minimize intertrial variability. For pharmacological experiments 10 µM gabazine and 5 µM 1-(2,4-Dichlorophenyl)-5-(4-iodophenyl)-4-methyl-N-1-piperidinyl-1H-pyrazole-3-carboxamide (AM251) were applied and were purchased from (Sigma-Aldrich). Membrane properties of human neurons did not show significant changes for up to 20 h after slicing, but recordings included in the analysis were arbitrarily terminated 15 h after slice preparation. Data were analyzed with Fitmaster (HEKA) and Origin 7.5 (OriginLab) Data are given as mean±standard deviation (S.D.). The Mann-Whitney U-test was used to compare datasets; differences were considered significant if p<0.05.

### Firing classification analysis

First, a set of n=200 electrophysiological features had been calculated for each cell identified based on light microscopic investigation of the axonal arbour. Then a wrapper feature selection method using Support Vector Machine (SVM) was used on the cells (rosehip cells: n=55; non-rosehip cells: n=52) to find the best feature set which separated the group of rosehip cells from the group of other cells. The best feature set consisted of 2 features, the maximal standard deviation of interspike-intervals (ISI SD) and the amplitude of sag in response to hyperpolarization (-100 pA). Sweeps were discarded with less than 5 spikes for the calculation of ISI SD. The sag value was calculated as the ratio of the voltages at the onset of the hyperpolarizing step and during steady state.

### Measurement of impedance profile

The impedance profile was determined by sinusoidal current injections using a standard exponential chirp pattern (0.2-200 Hz, 10 s duration) generated with Patchmaster (HEKA). Measurements (7-10 traces per cell) were made at resting membrane potential and the peak to peak amplitude of the command current waveform was tuned between 40-100 pA to test subthreshold voltage responses. The impedance profile (Z) was determined for each trace by calculating the fast Fourier transform (FFT) of the voltage response and dividing the FFT component of the corresponding command current, then the impedance profiles were normalized to the value at 200 Hz. After anatomical identification of the recorded cells, the dataset was pooled according to three defined cell types then the averaged impedance plotted against input frequency. For statistical comparison of the impedance profiles, four parameters were considered: impedance at lowest frequency (Z_0.2Hz_); cutoff frequency (f_cutoff_); resonance magnitude (Q, the impedance magnitude at the resonance peak i.e. maximal impedance value divided by the impedance magnitude at the lowest input frequency of 0.2 Hz); and the frequency at maximum impedance (f_max_).

### Two-photon calcium imaging

Structural labelling of rosehip cells was based on 40 µM Alexa Fluor 594 (Molecular Probes). We also applied 100 µM of Oregon Green 488 BAPTA-1 (Molecular Probes), in order to measure intracellular Ca^2+^ dynamics of pyramidal cell dendrites in the intracellular solution (see above). Imaging with multiphoton excitation was performed using a modified Zeiss LSM7 MP (Oberkochen, Germany) two-photon laser scanning system and a FemtoRose 100 TUN (R&D Ultrafast Lasers, Hungary) titanium–sapphire laser with Finesse4 pumping laser (Laser Quantum, UK) providing 100 fs pulses at 80 MHz at a wavelength of 820 nm. Fluorescence images were acquired through a x40 water-immersion objective (0.8 NA; Olympus, Japan).

### Single cell reverse transcription and digital PCR

At the end of electrophysiological recordings, the intracellular content was aspirated into the recording pipettes by application of a gentle negative pressure while maintaining the tight seal. Pipettes were then delicately removed to allow outside-out patch formation, and the content of the pipettes (~1.5 μl) was expelled into a low-adsorbtion test tube (Axygen) containing 0.5 μl SingleCellProtectTM (Avidin Ltd. Szeged, Hungary) solution in order to prevent nucleic acid degradation and to be compatible with direct reverse transcription reaction. Samples were snap-frozen in liquid nitrogen and stored or immediately used for reverse transcription. Reverse transcription of individual cells was carried out in two steps. The first step was performed for 5 min at 65°C in a total reaction volume of 7.5 μl containing the cell collected in 4 μl SingleCellProtect (Avidin Ltd., Cat.No.: SCP-250), 0.45 μl TaqMan Assays (Thermo Fisher), 0.45 μl 10 mM dNTPs (Thermo Fisher, Cat.No.: 10297018, 1.5 μl 5X first-strand buffer, 0.45 μl 0.1 mol/L DTT, 0.45 μl RNase inhibitor (Thermo Fisher, Cat.No.:N8080119) and 100 U of reverse transcriptase (Superscript III, Thermo Fisher, Cat.No.: 18080055). The second step of the reaction was carried out at 55°C for 1 hour and then the reaction was stopped by heating at 75°C for 15 min. The reverse transcription reaction mix was stored at −20°C until PCR amplification.

For digital PCR analysis, the reverse transcription reaction mixture (7.5 μl) was divided into two parts: 6 μl was used for amplification of the gene of interest and 1.5 μl cDNA was used for amplifying the housekeeping gene, GAPDH. Template cDNA was supplemented with nuclease free water to a final volume of 8 μl. 2 μl TaqMan Assays (Thermo Fisher), 10 μl OpenArray Digital PCR Master Mix (Thermo Fisher, Cat.No.: 4458095) and nuclease free water (3 μl) were mixed to obtain a total volume of 20 μl and the mixture was evenly distributed on 4 subarrays (256 nanocapillary holes) of an OpenArray plate by using the OpenArray autoloader. Processing of the OpenArray slide, cycling in the OpenArray NT cycler and data analysis were done as previously described ^84^. For our dPCR protocol amplification, reactions having CT values less than 23 or greater than 33 were considered primer dimers or background signals, respectively, and excluded from the data set.

### Histology and reconstruction

Following electrophysiological recordings, slices were immersed in a fixative containing 4% paraformaldehyde (for immunohistochemistry) or 4% paraformaldehyde, 15% (v/v) saturated picric acid and 1.25% glutaraldehyde (for reconstructions) in 0.1 M phosphate buffer (PB; pH=7.4) at 4˚C for at least 12 h. After several washes with 0.1 M PB, slices were frozen in liquid nitrogen then thawed in 0.1 M PB, embedded in 10% gelatin and further sectioned into 60 µm slices. Sections were incubated in a solution of conjugated avidin-biotin horseradish peroxidase (ABC; 1:100; Vector Labs) in Tris-buffered saline (TBS, pH=7.4) at 4˚C overnight. The enzyme reaction was revealed by 3’3-diaminobenzidine tetra-hydrochloride (0.05%) as chromogen and 0.01% H_2_O_2_ as oxidant.

Sections were post fixed with 1% OsO_4_ in 0.1 M PB. After several washes in distilled water, sections were stained in 1% uranyl acetate, dehydrated in ascending series of ethanol. Sections were infiltrated with epoxy resin (Durcupan) overnight and embedded on glass slides. Three dimensional light microscopic reconstructions were carried out using Neurolucida system (MicroBrightField) with 100x objective. Reconstructed neurons were quantitatively analyzed with NeuroExplorer software (MicroBrightField).

### Immunohistochemistry of biocytin-labeled cells

The recorded cells were first visualized with incubation in Cy3-conjugated streptavidin (Jackson Immunoresearch) for 2 h, diluted 1:400 in TBS. After examination by epifluorescence microscopy, the sections containing the soma of the labeled neurons were incubated in 20% normal horse serum in TBS to block nonspecific antibody-binding sites. Free-floating sections containing the soma were incubated in primary antibodies dissolved in TBS containing 0.05% NaN_3_ for 72 hours at room temperature. The following primary antibodies were used: rabbit-anti-pro-cholecystokinin (1:2000, gift from Andrea Varro, Liverpool University); mouse-anti-CNR1 (1:4000, ImmunoGenes); rabbit-anti-GABA (1:1000, Sigma-Aldrich); mouse-anti-NR2F2 (1:700, Abcam); mouse-anti-PV (1:1500, Swant); rabbit-anti-nNOS (1:1000, Cayman Chemical); rabbit-anti-NPY (1:300, Peninsula Laboratories); rat-anti-somatostatin (1:50, Merck Millipore); rabbit-anti-calbindin (1:2000, Swant); goat-anti-calretinin (1:700, Swant); goat-anti-acetyltransferase (1:100, Merck Millipore). After several washes in TBS, the immunoreactions were visualized with A488- or Cy5-conjugated secondary antibodies (1:500, Jackson Immunoresearch). The sections were mounted on slides in Vectashield (Vector Laboratories). Images were taken by confocal laser scanning microscope (LSM 880, Zeiss) using a 40x oil-immersion objective (1.4 NA). After photography, the sections were demounted, washed in 0.1 M PB, and biocytin was visualized with the avidin-biotinylated horseradish peroxidase method described above.

### Electron microscopy

Axonal boutons of biocytin filled rosehip (n=3) and neurogliaform cells (n=2) (identified based on distinctive electrophysiological properties and light microscopic investigation of the axonal arbour) were re-embedded and re-sectioned at 70 nm thickness. Digital images of serial EM sections were taken at magnifications ranging from 8,000x to 50,000x with a JEOL JEM-1400Plus electron microscope equipped with a 8 M pixel CCD camera (JEOL *Ruby*). Axon terminals were reconstructed in 3D and their volumes were measured using the Reconstruct software (<u>http://synapses.clm.utexas.edu/</u>) (n=31 boutons of rosehip cells; n=24 boutons of neurogliaform cells). The areas of active zones of rosehip cells were measured at perpendicularly cut synapses, where the rigid apposition of the pre- and postsynaptic membranes was visible (n=11 active zones).

## Acknowledgements

The authors thank the Allen Institute for Brain Science founders, Paul G. Allen and Jody Allen, for their vision, encouragement and support. This work was supported by the ERC Interimpact project (GT), the Hungarian Academy of Sciences (GT), the National Research, Development and Innovation Office of Hungary GINOP-2.3.2-15-2016-00018, VKSZ-14-1-2015-0155 and by the National Brain Research Program, Hungary (GT). The authors thank Lena Christiansen and Fan Zhang from Illumina, Inc. for their assistance with RNA sequencing.

## Author Contributions

Conceptualization, E.S.L., R.S.L., G.T.

Methodology, R.D.H., M.N., J.L.C., P.B., L.G.P., G.T. Validation, J.L.C., S.L.D., G.M., G.T.

Formal Analysis, T.B., B.D.A., J.M.M., J.A.M., P.V., M.R., S.B., R.H.S., E.B., J.B., G.O., G.M.

Investigation, R.D.H., M.N., J.L.C., F.D.F., S.S., K.A.S., A.W., D.N.T., E.B., J.B., A.K.K., N.F., B.K., M.R., G.M., A.O.,G.O., G.T.

Resources, E.S.L., F.J.S., N.J.S., R.H.S., R.S.L., G.T.

Data Curation, T.B., B.D.A., J.M.M., J.A.M., P.V., S.B.

Writing – Original Draft, T.B. R.D.H., J.L.C., J.A.M., E.S.L., E.B., G.O., G.T.

Writing – Review and Editing, T.B., R.D.H., J.A.M., R.H.S., R.S.L., E.S.L., G.T.

Visualization, T.B., R.D.H., J.L.C., S.L.D., J.A.M., E.B., G.M., G.O.

Supervision, E.S.L., F.J.S., N.J.S., R.H.S., R.S.L, G.T.

Project Administration, E.S.L., S.M.S., G.T.

Funding Acquisition, E.S.L., F.J.S., R.S.L., G.T.

## Competing Financial Interests

The authors declare no competing financial interest.

## Supplementary Information

**SI Figure 1.**
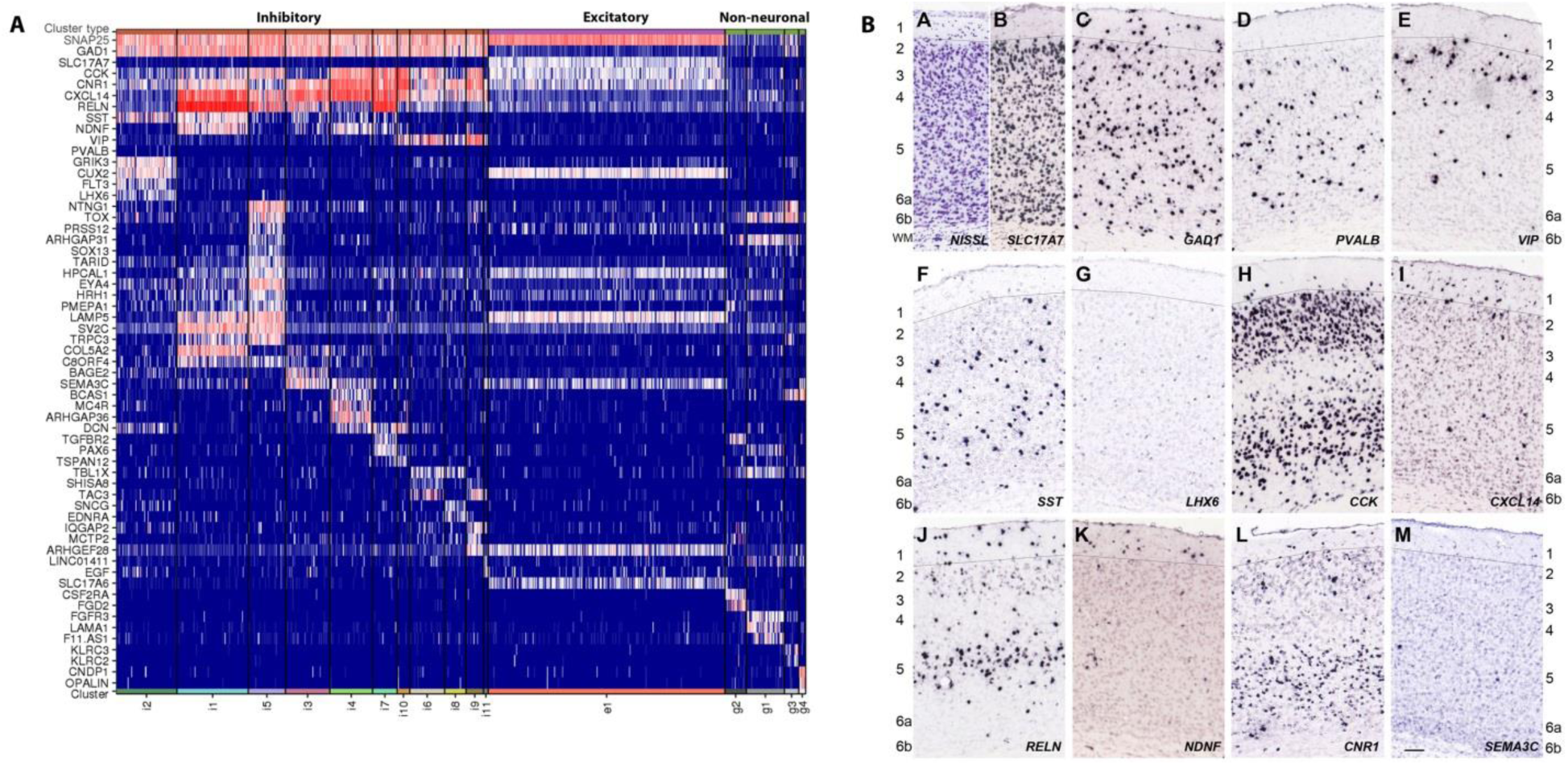
Marker gene expression patterns across identified single nuclei clusters. **A**, Heatmap of log-normalized expression (CPM) in single nuclei grouped by clusters that have been ordered by transcriptomic similarity. Canonical gene markers (*SNAP25*, *GAD1*, *SLC17A7*) classify clusters into broad classes of excitatory and inhibitory neuron and non-neuronal cell types. Within these broad types, many genes discretely mark individual clusters. **B**, Colorimetric ISH in mouse cortex of marker genes shown for human temporal cortex in Figure 1. Red arrows highlight cells with expression in layer 1.

**SI Figure 2.**
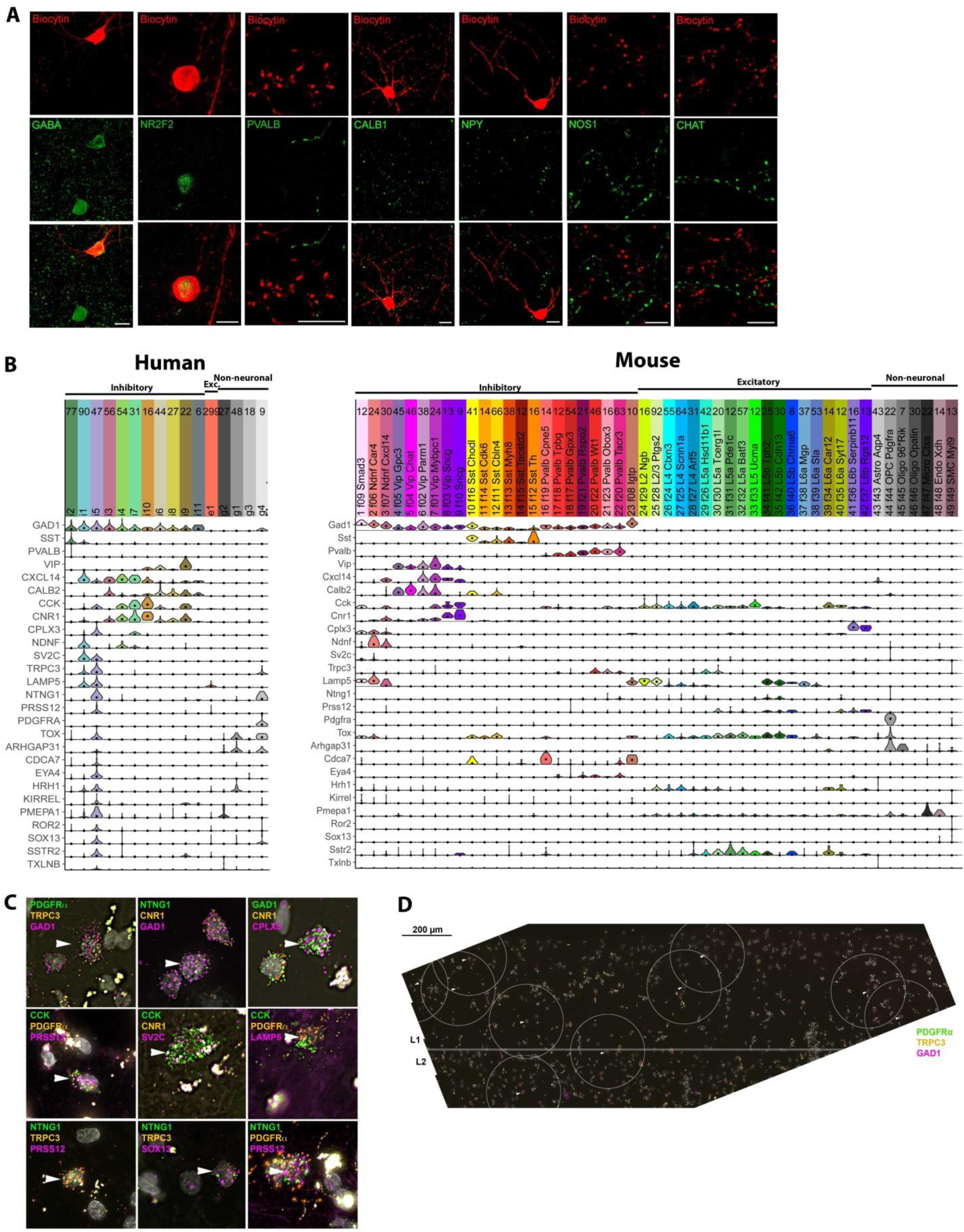
Additional molecular phenotype information of rosehip cells in layer 1 of the human cerebral cortex. **A**, Further immunolabeling showed that rosehip cells were immunopositive for gamma-aminobutyric acid (GABA) (n=2), and for chicken ovalbumin upstream promoter transcription factor II (COUP-TFII) (n=2). In addition, all tested rosehip cells were negative for many common interneuron markers including parvalbumin (n=3), neuronal nitric oxide synthase (n=4), neuropeptide Y (n=2), calbindin (n=2), and choline acetyltransferase (n=3). **B**, Expression distributions of marker genes in all cell types in human temporal cortex Layer 1 (left) and mouse primary visual cortex (right; data from ^4^). **C**, Multiplex FISH validation of rosehip marker co-expression. Arrowheads show examples of rosehip cells based on marker gene expression identified from RNA-Seq data. **D**, Multiplex FISH of approximately 2 mm section of Layer 1 and upper layer 2 with rosehip interneurons labeled based on marker gene expression. 300 µm diameter circles approximate the maximum extent of rosehip axonal arbors (see Figure 2E).

